# Structure of the TELO2-TTI1-TTI2 complex and its function in TOR recruitment to the R2TP chaperone

**DOI:** 10.1101/2020.11.09.374355

**Authors:** Mohinder Pal, Hugo Muñoz-Hernandez, Dennis Bjorklund, Lihong Zhou, Gianluca Degliesposti, J. Mark Skehel, Emma L. Hesketh, Rebecca F. Thompson, Laurence H. Pearl, Oscar Llorca, Chrisostomos Prodromou

## Abstract

The R2TP (RUVBL1-RUVBL2-RPAP3-PIH1D1) complex, in collaboration with HSP90, functions as a chaperone for the assembly and stability of protein complexes, including RNA polymerases, snRNPs and PI3 kinase-like kinases (PIKK) such as TOR and SMG1. PIKK stabilisation depends on an additional complex of TELO2, TTI1 and TTI2 (TTT), whose structure and function are poorly understood. We have now determined the cryo-EM structure of the human R2TP-TTT complex that together with biochemical experiments reveals the mechanism of TOR recruitment to the R2TP-TTT chaperone. The HEAT-repeat TTT complex binds the kinase domain of TOR, without blocking its activity, and delivers TOR to the R2TP chaperone. In addition, TTT regulates the R2TP chaperone by inhibiting RUVBL1-RUVBL2 ATPase activity and by modulating the conformation and interactions of the PIH1D1 and RPAP3 components of R2TP. Together, our results show how TTT couples the recruitment of TOR to R2TP with the regulation of this chaperone system.

## Introduction

The R2TP complex is a specialized co-chaperone of HSP90, consisting of an alternating heterohexameric ring of two AAA^+^ ATPases, RUVBL1 (also known as Pontin, TIP49, TIP49a and in yeast as Rvb1p) and RUVBL2 (also known as Reptin, TIP48, TIP49b and in yeast as Rvb2), a TPR-domain protein, RPAP3 (Tah1p in yeast) and a PIH-domain protein, PIH1D1 (Pih1p in yeast) (Martino et al., 2018; Rivera-Calzada et al., 2017). R2TP has been implicated in many biological processes, including the assembly, activation and stabilization of phosphatidylinositol 3 kinase-related kinases (PIKK) such as mTORC1, SMG1, ATM, DNA-PK, and ATR/ATRIP, the assembly of RNA polymerase II and snoRNPs, and in the assembly of axonemal dyneins (von Morgen et al., 2015). Most recently R2TP has been implicated in the function and assembly of replisome and nucleocapsids of RNA viruses responsible for measles and Ebola (Katoh et al., 2019; Morwitzer et al., 2019)

The structures of yeast and human R2TP complexes have been analysed by cryo-electron microscopy (cryoEM) (Martino et al., 2018; Rivera-Calzada et al., 2017; Tian et al., 2017). RUVBL1/Rvb1p and RUV/Rvb2p (henceforth R2) form a heterohexamer through interaction of the alternating AAA-domains of each subunit. Domain II (DII) from each subunit projects from the hexameric ring and defines the DII face of R2, while the AAA-ATPase domains define the opposing AAA-face. In all R2 complexes studied so far, other associated proteins predominantly bind to the DII-face of the ring, and in yeast and human R2TP complexes, the HSP90-recruiting TPR domains of RPAP3/Tah1p and the client-recruiting PIH1-containing PIH1/Pih1p (RPAP3/Tah1p-PIH1/Pih1p complex, henceforth TP) are located on this face of the ring. In human R2TP, RPAP3, a much larger protein than the yeast homologue Tah1p, contains a unique C-terminal □-helical domain that binds to the AAA-side of each RUVBL2 subunit. This RUVBL2-binding domain (RBD) provides a tight anchor for RPAP3 in complex with PIH1D1, while allowing for considerable flexibility of the TPR and N-terminal domains of RPAP3 at the DII-face of R2. Since each R2TP complex contains three copies of RUVBL2, up to three RPAP3-PIH1D1 complexes can bind per R2TP, but although flexibly tethered, only one PIH1D1 at a time can interact directly to R2. One molecule of PIH1D1 binds to the DII domain of one RUVBL2 subunit, inducing conformational changes that facilitate nucleotide exchange in this subunit and promote ATP turnover (Munoz-Hernandez et al., 2019).

Recruitment of PIKK proteins to the R2TP complex requires an additional adaptor complex formed by TELO2 (Takai et al., 2007) and its associated proteins TTI1 and TTI2 (Takai et al., 2010) known as TTT. Recruitment of TTT to R2TP is believed to be mediated through interaction of a CK2-phosphorylated acidic motif in the flexible linker connecting the N- and C-terminal domains of TELO2, with the PIH domain of PIH1D1 (Horejsi et al., 2014; Horejsi et al., 2010; Pal et al., 2014). TELO2 interaction with PIKKs has itself been suggested to be dependent on HSP90, although the mechanistic basis for this has not been defined (Takai et al., 2010).

We show that contrary to expectations TTT can be directly recruited to R2 and we have determined the structure of the human R2-TTT complex by single-particle cryo-EM. The structure permits atomic modelling of the RUVBL1 and RUVBL2 and tracing of the α-helical repeats that form the solenoid structures of the TTT proteins. Our data reveals a direct engagement of the RUVBL1-RUVBL2 hetero-hexamer by the TTT complex that is independent of the RPAP3-PIH1D1 (TP) components. TTT interacts with two DII domains from consecutive RUVBL1 and RUVBL2 subunits and this constrains the conformational flexibility of the DII domains of the RUVBL proteins in the ADP-bound state, inhibits the conformationally-coupled activity of the RUVBL AAA+-ATPase, and disturbs the engagement of PIH1D1 to the DII domains. We show that TTT is fully competent to bind a strongly R2TP-dependent PIKK client, TOR, but does so without inhibiting TOR kinase activity. Together, our cryo-EM structures and biochemical experiments reveal that TTT couples the recruitment of TOR to the R2TP chaperone with the regulation of the chaperone.

## Results

### Cryo-EM of the RUVBL1-RUVBL2-TTT complex

Although, current models suggest that TTT recruitment to R2TP is mediated by the specific interaction of PIH1D1 with a CK2 phosphorylation site on TELO2 (Horejsi et al., 2014), we found that we could readily pull-down complexes of TELO2-TTI1-TTI2 with strep-tagged RUVBL1-RUVBL2 (R2-TTT) in the absence of RPAP3 and PIH1D1 **(Fig. 1A)** and that R2 and TTT co-migrated in size exclusion chromatography as a stable complex **(Fig. S1A-C)**. The TTI1-TTI2 subcomplex (henceforth TT) was also co-precipitated by RUVBL1-RUVBL2 (R2-TT), demonstrating that TELO2 was also not required for complex formation **(Fig. 1A)**.

**Fig. 1.**
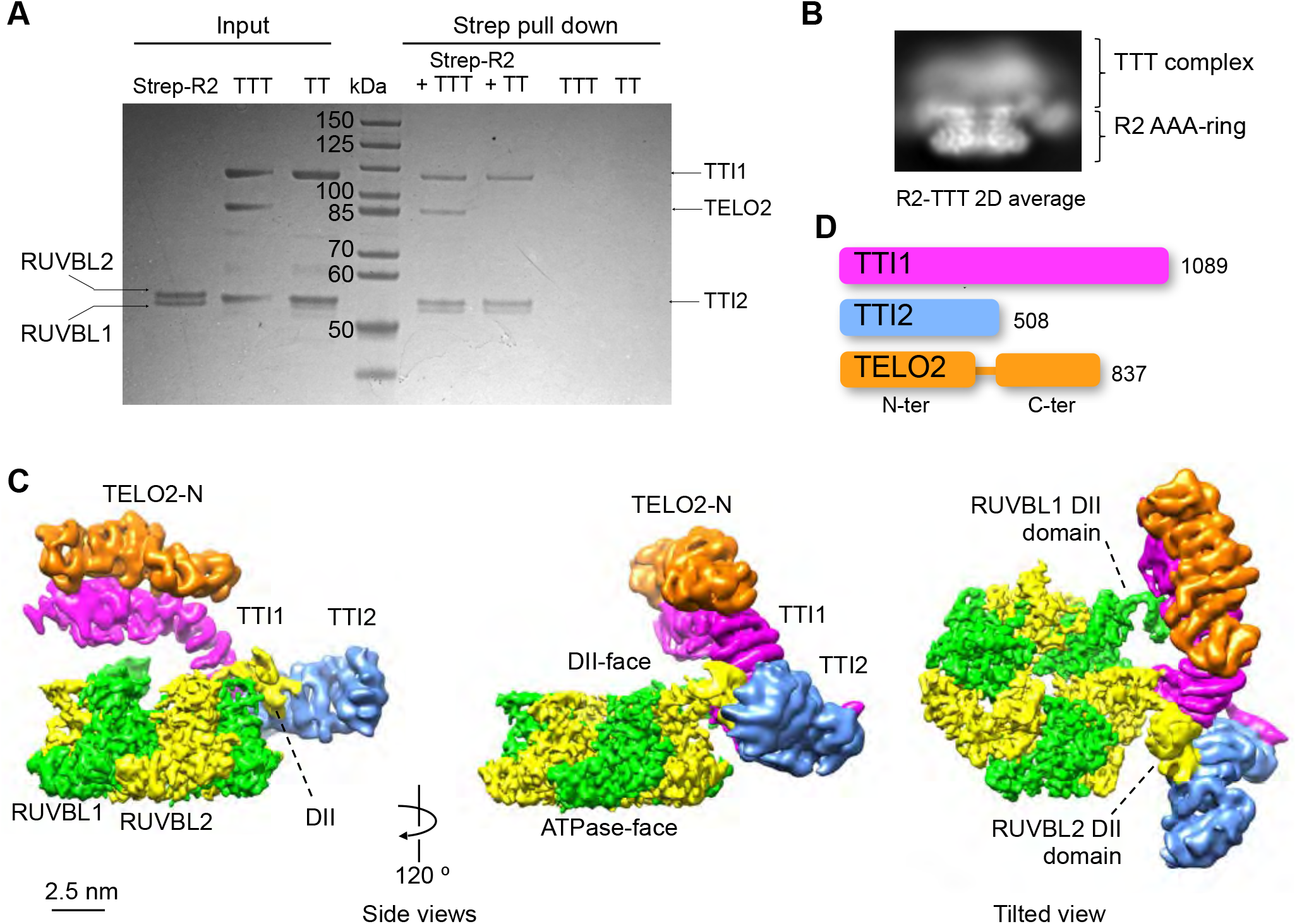
Cryo-EM of the human RUVBL1-RUVBL2-TTT complex. **A.** Coomassie-stained SDS-PAGE gel showing analysis of interaction of RUVBL1-RUVBL2 (R2), TTI1-TTI2-TELO2 (TTT) and TTI1-TTI2 (TT) sub-complexes. Pulldown on the strep-tag attached to the C-terminus of RUVBL2 co-precipitates TTT and TT sub-complexes, showing that neither RPAP3, PIH1D1 nor TELO2 are required to couple TTI1-TTI2 to R2. TTT and TT are not pulled down in the absence of R2. **B.** Representative 2D average of R2-TTT cryo-EM data. **C.** Several views of human R2-TTT volume. RUVBL1-RUVBL2 hexamer – green and yellow respectively. Three distinct α-helical segments (TTI1 - magenta, TTI2 - blue and TELO2-N - orange) are identifiable in the cryo-EM reconstruction after multi-body refinement of the R2-TTT complex. **D.** Schematic showing the relative lengths of the TTT components TTI1, TTI2 and TELO2. All three proteins are known or believed to be entirely composed of HEAT repeats, which is interrupted by a ~70 residue flexible linker in TELO2, that connects the N- and C-terminal domains.

The assembled R2-TTT complex was vitrified by plunging into liquid ethane and Titan Krios microscopes equipped with Gatan K2 detectors were used to collect cryo-EM data. 2D averages showed that TTT occupies the DII-face of the RUVBL1-RUVBL2 ring **(Fig. 1B)**. First observations during 3D refinement of the cryo-EM data revealed that TTI1-TTI2-TELO2 binds to the DII domains of consecutive RUVBL1 and RUVBL2 subunits, and that we could saturate each R2 ring with up to 3 TTTs by increasing the TTT vs R2 ratio during incubation. We exploited the presence of several copies of TTT per particle making use of a symmetry expansion strategy **(see METHODS and Fig. S2A)**. We also observed that TTT attaches flexibly to the DII domains and we split the volume in two bodies. One body contained the ATPase ring of R2 without the flexible DII domains and this refined to a resolution of 3.41Å. The second body included TTT and the two DII domains making direct contact with TTT and this reached an average resolution of 5.02Å **(Fig. S2C)**. Then we built a composite map combining both bodies **(Fig. 1C)**.

In the cryo-EM map we could readily identify the heterohexameric ring formed by RUVBL1 and RUVBL2 but in addition, there is a strong body of density at the edge of the ring, projecting from the face that bears the RUVBL-DII domains, corresponding to TTT (**Fig. 1C**). TELO2, TTI1 and TTI2 are known or predicted to contain substantial regions of HEAT repeat (Kaizuka et al., 2010; Takai et al., 2010) and the density for TTT appears as a large curved arc containing more than 30 α-helices in a solenoid arrangement which becomes increasingly poorly defined at one end (magenta colour); a smaller segment of ~20 α-helices that lies on top of the large segment towards the less well-ordered end (orange colour); and a second shorter curved arc in which at least 15 helices can be identified, which abuts the well-ordered end of the large arc (blue colour) **(Fig. 1C**).

Based on its size (**Fig. 1D**), we identify the larger segment as TTI1, however the attribution of the two smaller segments as TTI2 or TELO2 is less clear. We identified TELO2 within the three segments of helical solenoids by obtaining a reconstruction from an R2-TT pull-down complex the lacks TELO2. While this was only of limited resolution (~8Å), density corresponding to much of the helical solenoids was evident **(Fig. S3)**. However, when the R2-TT map was compared with that for the R2-TTT complex, the segment in orange colour was not observed, even at a very low contour level, suggesting that it is due to the absence of TELO2 in the R2-TT complex. R2 does not contact TELO2 directly, which agrees TELO2 being dispensable for the interaction of TT with R2 **(Fig. 1A)**.

### Atomic model of human TTT complex

We built an atomic model of TTT by assigning of regions of HEAT repeat in the cryo-EM map and resolving uncertainties by mapping interactions between truncated versions of the TTT components. The crystal structure of an engineered construct of yeast Tel2p (Takai et al., 2010) consists of two helical solenoid segments of 19 and 9 helices respectively, separated by ~60 unstructured residues that include the phosphorylated binding motif for the PIH domain of Pih1p (Horejsi et al., 2014; Pal et al., 2014). The size and shape of the segment assigned to TELO2 in the cryo-EM structure **(Fig. 1C**, orange colour) corresponds well with the structure of the larger N-terminal segment of yeast Tel2p, and a homologous model of the corresponding region of human TELO2 could be readily docked into this density in the R2-TTT map as a rigid body **(Fig. 2A)**. The substantial interface between TELO2 N-terminal domain (TELO2-N) and TTI1 is consistent with the identification of the N-terminal segment of Tel2p rather than the C-terminal segment as the main interactor with Tti1p-Tti2p (Takai et al., 2010). Based on this we identified an additional region of density at a lower contour level, adjacent to C-terminal end of TELO2 N-terminal domain **(Fig. 2B)**. This likely corresponds to the C-terminal domain of TELO2 (TELO2-C) which is known to connect to the N-terminal domain by a linker that would allow the flexible tether of TELO2-C to the rest of TTT **(Fig. 2B)**. Based on this assignment, we can then attribute the remaining segment to the ~500 residues of TTI2.

**Fig. 2.**
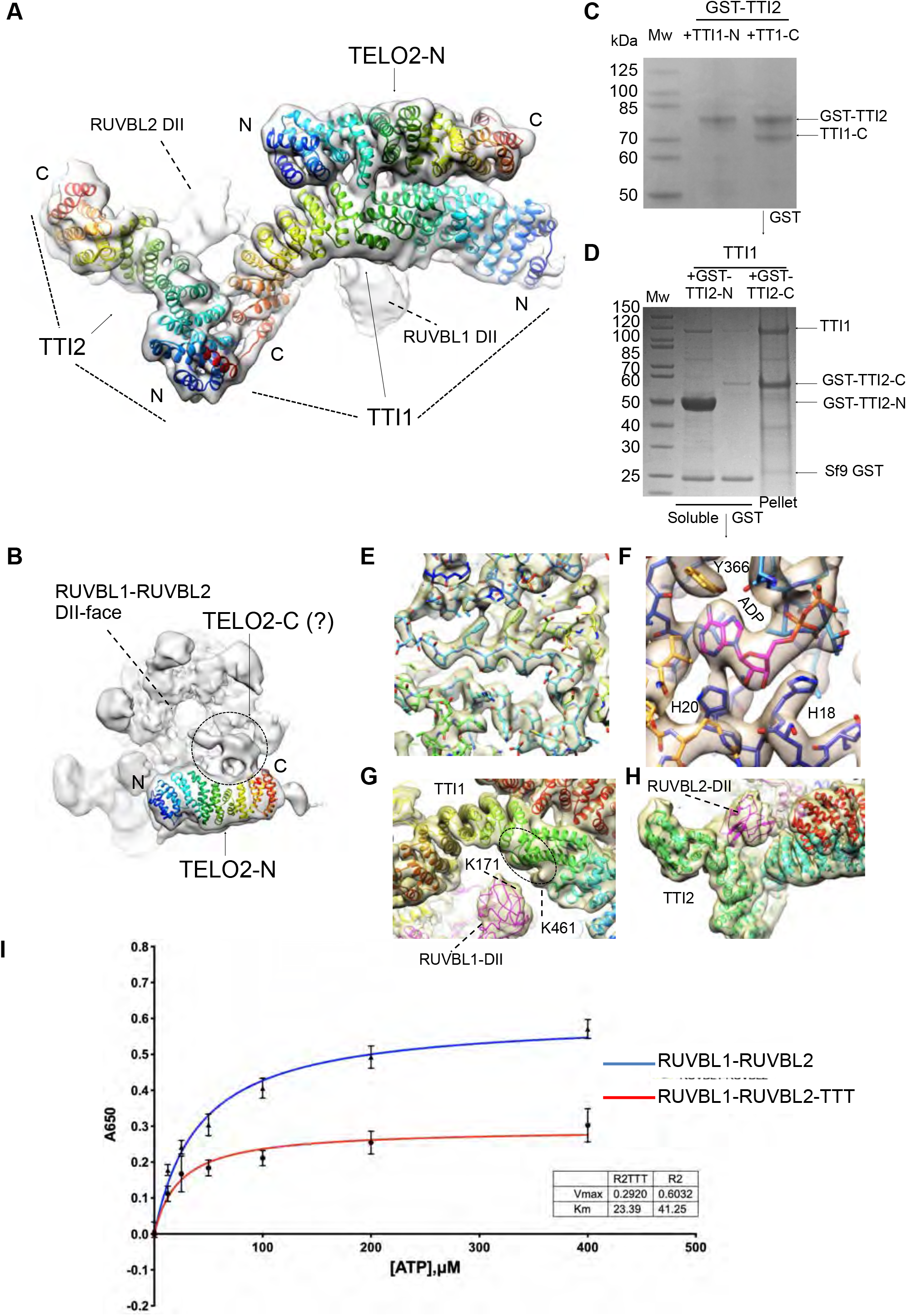
Molecular modelling of TTT and its interaction with RUVBL1-RUVBL2. **A.** Molecular model of TTT fitted within the cryo-EM density of R2-TTT (transparent density). Individual helices of TTI1 and TTI2 were fitted manually into density using COOT, and connected into a continuous structure by taking advantage of the repetitious linear right-handed α-solenoid architecture of HEAT repeats. The polarity of the two polypeptides was determined by the results of the interaction mapping experiments shown in **C** and **D**. The fit of the poly-alanine structures was optimised by real-space refinement in Phenix (*18*). A homology model of the N-terminal domain of human TELO2 based on the crystal structure of *S.cerevisiae* Tel2p (*9*), was manually adjusted to fit the experimental density using COOT (*17*). The TTT model was optimised by real-space refinement in Phenix (*18*). Each protein is coloured as a rainbow from N- to C-terminal ends. **B.** A density in the R2-TTT map projects from the C-terminal end of TELO2-N and which is sufficient to accommodate an homology model of the structured region of the C-terminal domain of human TELO2 (not shown). The gap between the docked TELO2 N- and C-terminal domains could be comfortably spanned by the ~70 residue intrinsically disordered linker segment that connects them in the primary structure. **C.** Co-expression of GST-tagged full-length TTI2 with TTI1 constructs spanning residues 1-459 (TTI1-N) and residues 468-1089, in Sf9 insect cells. Only the TTI1-C construct was co-precipitated as a soluble species with GST-TTI2, suggesting that TTI2 primarily interacts with the C-terminal half of TTI1. **D.** Co-expression of full-length TTI1 with GST-tagged TTI2 constructs spanning residues 1-193 (TTI2-N) and residues 194-508 (TTI2-C), in Sf9 insect cells. The GST-tagged TTI2-N was well expressed as a soluble species and co-precipitated TTI1 in GST pull-downs, whereas co-expressed TTI1 and GST-TTI2-C were primarily found in the insoluble fraction. This suggests that TTI1 primarily interacts with the NN-terminal half of TTI2. **E.** Close up of electron density in the core of the RUVBL2 subunit that contacts TTI1 and TTI2, showing clear electron density for side chains. The fit of the all-atom model of the RUVBL1-RUVBL2 heterohexamer was optimised by real-space refinement in Phenix. **F.** Close-up of electron density in the nucleotide-binding site of the RUVBL1 subunit that contacts TTI1, showing bound ADP. All six nucleotide-binding sites in the R2 ring are fully occupied by ADP, and the His-Ser-His motif that closes off the nucleotide-binding site as part of the N-terminal ‘gatekeeper’ segment (*7*), is also fully ordered in all six ATPase subunits. **G.** Interaction of TTI1 (rainbow coloured helices) with the DII domain of a RUVBL1 subunit (magenta). The fit of the all-atom model of the RUVBL1-RUVBL2 heterohexamer and the TELO2-N domain, and the poly-Ala models of TTI1 and TTI2 were simultaneously optimised by real-space refinement in Phenix against a ‘consensus’ electron density map comprising maps from separate multibody and focussed refinements of the R2 rings and TTT with the interacting DII domains. In an XL-MS analysis of R2TP-TTT a single cross-link was observed between RUVBL1-Lys171 (black circle) and Lys461 of TTI1. Although the resolution of the TTI1 density doesn’t permit fitting of the sequence, the crosslink helps localise the approximate location of TTI1-Lys461 (red ellipse). **H.** Interaction of TTI1 (cyan helices) and TTI2 (green helices) with the DII domain of a RUVBL2 subunit. The DII domain packs into the cleft formed at the junction of the α-helical solenoids of the two HEAT repeat proteins. **I.** Human RUVBL1-RUVBL2 displays a weak inherent ATPase activity with a maximal rate of ~0.6 mole/min/mole. This is significantly diminished in the presence of TTT at a 3:1 molar ratio.

No prior structures are available for TTI1 or TTI2 or homologues thereof, intact or in part. While the local resolution of the maps in these regions (4.5 – 5Å) is sufficient to define the positions and topological connection of the α-helices making up the HEAT repeats that constitute the majority of these two proteins, it is not high enough to allow assignment of amino sequence or the orientation of the Cα-Cβ vectors of the side chains which indicates the N→C polarity of the helix. Consequently, the direction of the polypeptide chain in the α-helical solenoid cannot be directly determined.

To solve this problem, we took advantage of the effectively linear mapping of primary to tertiary structure in α-helical solenoid architectures such as the HEAT repeats that are predicted to form the majority of TTI1 and TTI2. One end of the helical density ascribed to TTI2 comes into close contact with one end of the long arc of helical density ascribed to TTI1. To determine which end of the TTI1 sequence this corresponds to, we co-expressed separate strep-tagged N-terminal (1-459) and C-terminal (468-1089) parts of TTI1 with glutathione S-transferase (GST) tagged-TTI2FL in insect cells. Strep-TTI1-C was clearly recovered along with GST-TTI2 in glutathione bead pull-downs from TTI1-C expressing cells, whereas no strep-TTI1-N was observed in similar experiments with the TTI1-N expressing cells **(Fig. 2C)**. Correspondingly GST-TTI2 was co-precipitated with strep-TTI1-C, but neither strep-TTI1-N nor GST-TTI2 were recovered by strep-tag pull-down from strep-TTI-N expressing cells **(Fig. S4)**. In a similar experiment we coexpressed full length TTI1 in insect cells, with either the N-terminal (1-193) or C-terminal (194-508) part of TTI2 as a C-terminal fusion to GST. We found that the GST-TTI2-N was highly expressed as a soluble protein, and co-purified full-length untagged TTI1 in glutathione-bead pull-downs. In contrast, the GST-TTI2-C fusion was barely detectable in the soluble fraction and did not co-precipitate TTI1 **(Fig. 2D).** Taken together these results indicate that the TTI1-TTI2 interface involves the N-terminus of TTI2 binding to the C-terminus of TTI1 with the more central / N-terminal part of TTI1 providing the binding site(s) for TELO2 N-terminal domain **(Fig. 2A)**. We were then able to dock known structures or homology models of RUVBL1, RUVBL2, TELO2-N and the helical repeats of TTI1 and TTI2 into the cryoEM maps, and optimize the atomic fits using restrained real-space refinement in phenix.refine (Adams et al., 2010) and manual adjustment in COOT (Casanal et al., 2019).

### RUVBL1-RUVBL2 – TTT Interactions regulate ATPase activity

The structure of the RUVBL1-RUVBL2 ring (maximum resolution 3.41Å) within the overall human complex, is similar to that previously described in the R2TP complex (Martino et al., 2018; Munoz-Hernandez et al., 2019).The amino acid sequence of all six subunits is well resolved with clear side chain density evident for most of the polypeptide chain (**Fig. 2E**), apart from the DII insertion domains. Two of these, which interact with the TTT proteins are resolved at resolutions in the 4.5 – 7.5Å range, while the other four are substantially disordered. All six subunits have clear density for bound ADP, with the N-terminal ‘gatekeeper’ segment of polypeptide incorporating the His-Ser-His motif, well-ordered and interacting with the bound nucleotide (Munoz-Hernandez et al., 2019) (**Fig. 2F**).

Although recruitment of TELO2 (Tel2p in yeast), and by presumption the entire TTT subcomplex, to R2TP was believed to depend on interaction of a conserved CK2 phosphorylation site in TELO2/Tel2p with the PIH domain of PIH1D1/Pih1p (Horejsi et al., 2014; Horejsi et al., 2010; Pal et al., 2014), our 3D structures and supporting biochemistry clearly show substantial direct interactions between elements of the TTT sub-complex and the RUVBL ring that are completely independent of the TP components – RPAP3/Tah1p and PIH1D1/Pih1p and are sufficient to form stable R2-TTT and R2-TT complexes.

At the heart of the TTT interface with R2, the concave inner surface of TTI1 wraps around the projecting DII domain of a RUVBL1 subunit, which contacts helices from the central part of the TTI1 solenoid, on the face opposite to the binding site of the TELO2-N segment (**Fig. 2G**). Consistent with this arrangement, we identified a single cross-link between RUVBL1 and TTI1 in a cross-linking mass spectrometry (XL-MS) analysis of a putative human R2TP-TTT complex (**Fig. S4**), connecting Lys 171 of RUVBL1 with Lys 461 of TTI1. While the resolution of the TTI1 density does not allow this residue to be pinpointed with any precision, it does localise it to a region encompassing one or two helical segments in the fitted model and helps establish the approximate staging of the sequence along that model. Simultaneously, the DII domain of the clockwise adjacent RUVBL2 subunit packs into the V-shaped cleft formed at the junction of the predicted C-terminus of TTI1 and the predicted N-terminus of TTI2 (see above), interacting with helices from the convex faces of both proteins (**Fig. 2H**). As essentially all the contacts between the TTT sub-complex and the R2 ring are mediated by the flexible DII domains, there is substantial conformational variability in the relative orientation of the TTT and R2 components as revealed by multi-body refinement (Nakane et al., 2018).

We previously showed a direct coupling between the conformation of a N-terminal region in RUVBL2 containing the His-Ser-His motif that binds nucleotide and the conformation of DII domains, which was promoted by their interaction with PIH1D1 (Munoz-Hernandez et al., 2019). This was shown to affect the binding of nucleotides to regulate nucleotide exchange as part of the AAA^+^ ATPase cycle. As the TTT complex makes extensive interaction with the DII domains of a pair of consecutive RUVBL1 and RUVBL2 proteins in the heterohexameric ring, we explored if TTT binding influenced the AAA^+^ ATPase cycle. Using a sensitive Malachite Green phosphate release assay, we were able to measure a basal ATPase activity with Vmax ~ 0.6 mole/min/mole for human RUVBL1-RUVBL2, which was substantially diminished by addition of a 3:1 molar equivalent of TTT **(Fig. 2I)**. Therefore, TTT directly binds RUVBL and modulates their ATPase activity.

### PIH1D1/Pih1p and TTT compete for the interaction with RUVBL1-RUVBL2

TTT is assumed to function in the context of R2TP, a complex where R2 also binds RPAP3-PIH1D1 (TP). As we have previously shown that PIH1D1 also interacts with the DII domains of R2 we explored whether TTT and PIH1D1 compete for interaction with R2 or can be accomodated together within a super-complex. For this, we assembled the human R2TP-TTT complex for cryo-EM studies by mixing separately purified R2TP and TTT complexes and the mix was analysed by cryo-EM.

We obtained a reconstruction for the R2TP-TTT complex with an average resolution of 6.1Å in which, as well as a bound TTT sub-complex, we could readily identify two well-defined copies of the C-terminal RUVBL2-binding domain (RBD) of RPAP3 bound to the AAA-face of the RUVBL1-RUVBL2 ring (Martino et al., 2018; Maurizy et al., 2018). (**Fig. 3A, Fig. S5**). As with previous studies a third RBD was evident at a lower contour level. The structural organization of TTT in R2TP-TTT was essentially identical to that observed in the R2-TTT complex. In R2TP, whereas three RPAP3-PIH1D1 complexes can be tethered to each RUVBL1-RUVBL2 ring at a time due to the interaction of the RPAP3-RBD domain to each of the three RUVBL2 subunits, only 1 PIH1D1 is effectively engaged to the DII domains (Martino et al., 2018; Munoz-Hernandez et al., 2019). Interestingly, although present in the *in vitro* assembled R2TP-TTT complex, no density was evident for PIH1D1 at the binding site identified in previous studies, nor for the conformationally flexible N-terminal regions of RPAP3 (Martino et al., 2018; Munoz-Hernandez et al., 2019). This suggests that TTT and PIH1D1 might compete for the binding to the DII domains of R2.

**Fig. 3.**
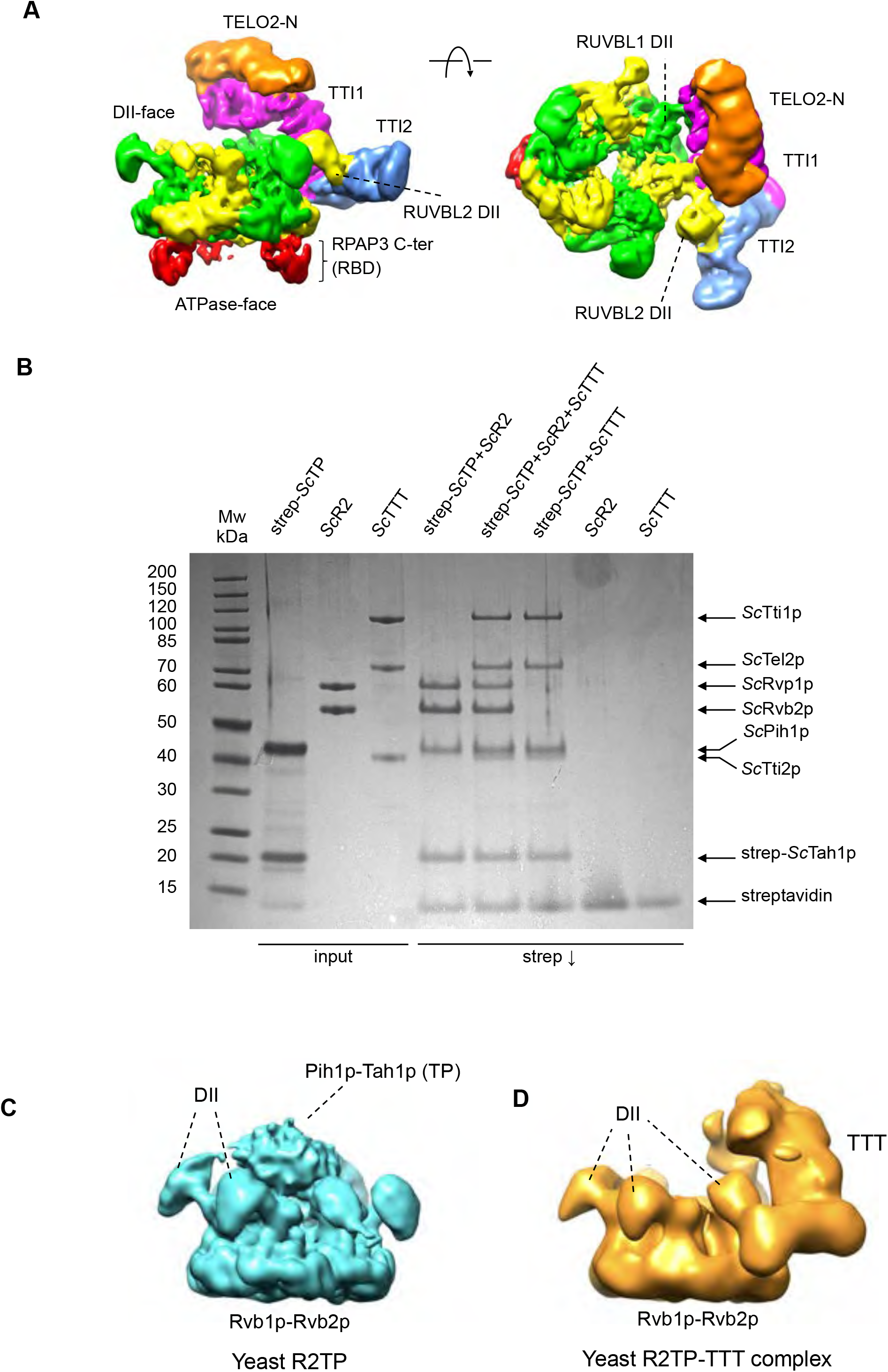
Cryo-EM of human and yeast R2TP-TTT complexes. **A.** Two views of human R2TP-TTT volume: RUVBL1-RUVBL2 hexamer – green and yellow respectively; TTI1 – magenta; TTI2 – blue; TELO2-N – orange. RPAP3 C-terminal RUVBL2-binding domain (RBD) is identifiable in the ATPase face of each of the RUVBL2 subunits – red. **B.** Coomassie-stained SDS-PAGE gel showing analysis of interaction of yeast Tah1p-Pih1p (yTP), Rvb1p-Rvb2p (yR2), Tel2p-Tti1p-Tti2p (yTTT) sub-complexes. Pull-down on the tandem-strep tag attached to the N-terminus of Tah1p within the yTP complex, coprecipitates yR2 and yTTT simultaneously and as separate co-complexes. **C.** Cryo-EM analysis of co-precipitated Tah1p-Pih1p-Rvb1p-Rvb2p-Tel2p-Tti1p-Tti2p (yeast R2TP-TTT). 3D classification yields two main classes – one class contained particles with strong central density between the DII insertion domains of the R2 ring comprising ~2/3rds of the particles, resembled the previously described yeast R2TP complex (EMD-3678) (Rivera-Calzada et al., 2017), **D.** Cryo-EM analysis of co-precipitated Tah1p-Pih1p-Rvb1p-Rvb2p-Tel2p-Tti1p-Tti2p (yeast R2TP-TTT). A second class contained particles with strong peripheral density at one edge of the R2 ring corresponding to the TTT complex. No significant class of particles was identified in which the central and peripheral density features were simultaneously present.

To further support this hypothesis, we also assembled an *S.cerevisiae* R2TP-TTT complex *in vitro*, by mixing a pre-assembled Rvb1p-Rvb2p-Pih1p-Tah1p complex with an equal concentration of a pre-assembled Tel2p-Tti1p-Tti2p complex where the components were purified from insect cells to ensure substantial post-translational modifications on the Tel2p component. Yeast TTT prepared in this way was co-precipitated in pull-down experiments with Tah1p-Pih1p (**Fig. 3B**), suggesting that any phosphorylation required within the Tel2p motif predicted to bind the PIH domain of Pih1p were present on the sample. Cryo-EM data were collected and processed as above. 3D classification of particles from this sample revealed a more heterogeneous distribution than for the human samples, which ultimately partitioned into two main classes. The more populated and higher resolution (~8Å) class possessed a strong centrally positioned feature nestled between the DII domains of the R2 ring, but with no density corresponding to TTT, and strongly resembled the previously described yeast Rvb1p-Rvb2p-Pih1p-Tah1p complex structure (Rivera-Calzada et al., 2017) (**Fig. 3C**). The second, less populated and lower resolution class (~13Å), lacked the central feature, but possessed a substantial additional density at the periphery of the R2 ring, which resembled the human TTT density in the human R2TTT reconstruction, and in which similar segments could be identified (**Fig. 3D**). No significant classes were found in 3D classification in which the central and peripheral density features were simultaneously present.

Therefore, our analysis of human and yeast R2TP bound to TTT suggest that TTT and PIH1D1 compete for their interaction with the DII domains.

### TTT delivers TOR to the R2TP chaperone

While our data suggest that TTT also plays a role in regulating the ATPase activity of the R2 ring, its main function is believed to be as an adaptor, recruiting PIKK client proteins such as TOR to the R2TP or R2 complex (Houry et al., 2018; Kakihara and Houry, 2012; Munoz-Hernandez et al., 2018). To test this, we purified *S.cerevisiae* TTT in which Tti1p carried a C-terminal flag-tag, and looked at its ability to interact with a TOR1-Lst8 complex from the closely related yeast *Kluveromyces marxianus* (construct a kind gift of Roger Williams, MRC-LMB, Cambridge) which has 83% sequence similarity to *S.cerevisiae* TOR2, but is far more amenable to large scale expression and purification. We found that KmTOR was co-immunoprecipitated by beads carrying an anti-flag antibody, in the presence of flag-tagged TTT, but not in its absence (**Fig. 4A**). Interestingly, KmLst8, which was evident in the input TOR1 preparation as a weak-staining band, was not visible in the immunoprecipitated material. Flag-tagged TTT was also able to simultaneously co-immunoprecipitate KmTOR and R2 when both components were present, suggesting that KmTOR, R2 and TTT can form a complex (**Fig. S6**). To determine which component(s) of the TTT complex are required for interaction with the PIKK client, we separately expressed Tel2p and the constitutive Tti1p-Tti2p complex and looked at their interaction with KmTOR. We found that KmTOR was effectively co-precipitated with Tti1p - GST-Tti2p in a GST pull-down experiment, but not by GST-Tel2p (**Fig. 4B**). In contrast to TTT, TP (Tah1p-Pih1p) or R2TP complexes in which Tah1p carried a tandem strep-tag, were unable to co-precipitate KmTOR directly. However, strep-tagged TP could co-precipitate KmTOR when TTT was also present, demonstrating that TTT can provide a bridging interaction between the two, in which the binding sites for TP and KmTOR on TTT are non-overlapping (**Fig. 4C**).

**Fig. 4.**
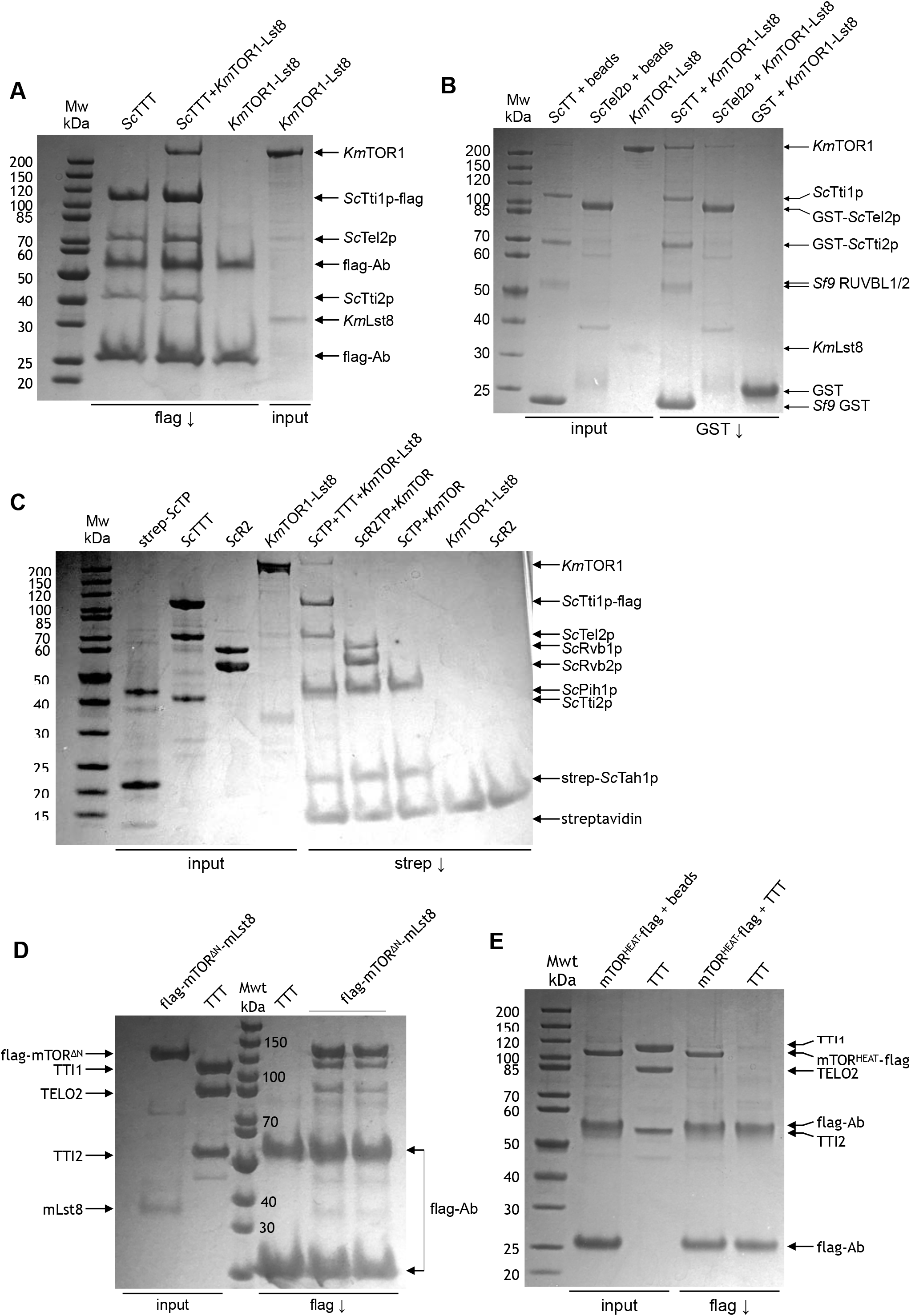
Mapping the interactions between TTT and TOR. **A.** Coomassie-stained SDS-PAGE gel showing interaction of *Kluveromyces marxianus* TOR1-Lst8 with *S.cerevisiae* TTT. Flag-tagged Tti1p, co-immunprecipates TOR1, whereas anti-flag beads alone do not. Lst8, which is evident as a weakly staining band in the input, was not evident in the immunoprecipitated TTT-TOR1. **B.** As **A**, but for interaction of KmTOR-Lst8 with GST-tagged Tti2p-Tti1p or GST tagged Tel2p. KmTOR was co-precipitated in GST pull-down with Tti1p-GST-TTI2p, but not with Tel2p. **C.** As **A**, but for interaction of KmTOR-Lst8 with TP and R2TP complexes containing strep-tagged Tah1p. Neither TP nor R2TP complexes were able to co-precipitate KmTOR directly. However, tagged TP could co-precipitate KmTOR when TTT was present, confirming that TP and KmTOR do not compete for binding to TTT. **D.** As **A**, but for interaction of a flag-tagged N-terminally truncated mTOR construct (mTOR^ΔN^) co-expressed with mLST8, with human TTT. TTT is co-immunoprecipitated by flag-tagged mTOR^ΔN^ but not by flag beads alone. **E.** As **A**, but for interaction of a flag-tagged soluble N-terminal fragment of mTOR encompassing the major segment of HEAT repeats (mTOR^HEAT^) with human TTT. Flag-tagged mTOR^HEAT^ was precipitated by flag beads, but did not co-immunoprecipitate TTT.

We also expressed and purified a flag-tagged construct of human mTOR (mTOR^ΔN^) which lacks the majority of the N-terminus (residues 1-1376), but retains the binding site for mLst8, and displays essentially full kinase activity (Yang et al., 2013).We found that human TTT complex was co-immunoprecipitated by beads carrying an anti-flag antibody, in the presence of a flag-tagged mTOR^ΔN^-mLst8 complex, but not in its absence (**Fig. 4C**). We also expressed and purified a soluble flag-tagged segment of mTOR comprising the first 929 residues. We found that this construct, which encompasses the main HEAT repeat segment of mTOR, was not able to co-precipitate human TTT (**Fig. 4E**), however this fragment was able to interact with RUVBL1-RUVBL2 in the absence of TTT (**Fig. 5A**).

**Fig. 5.**
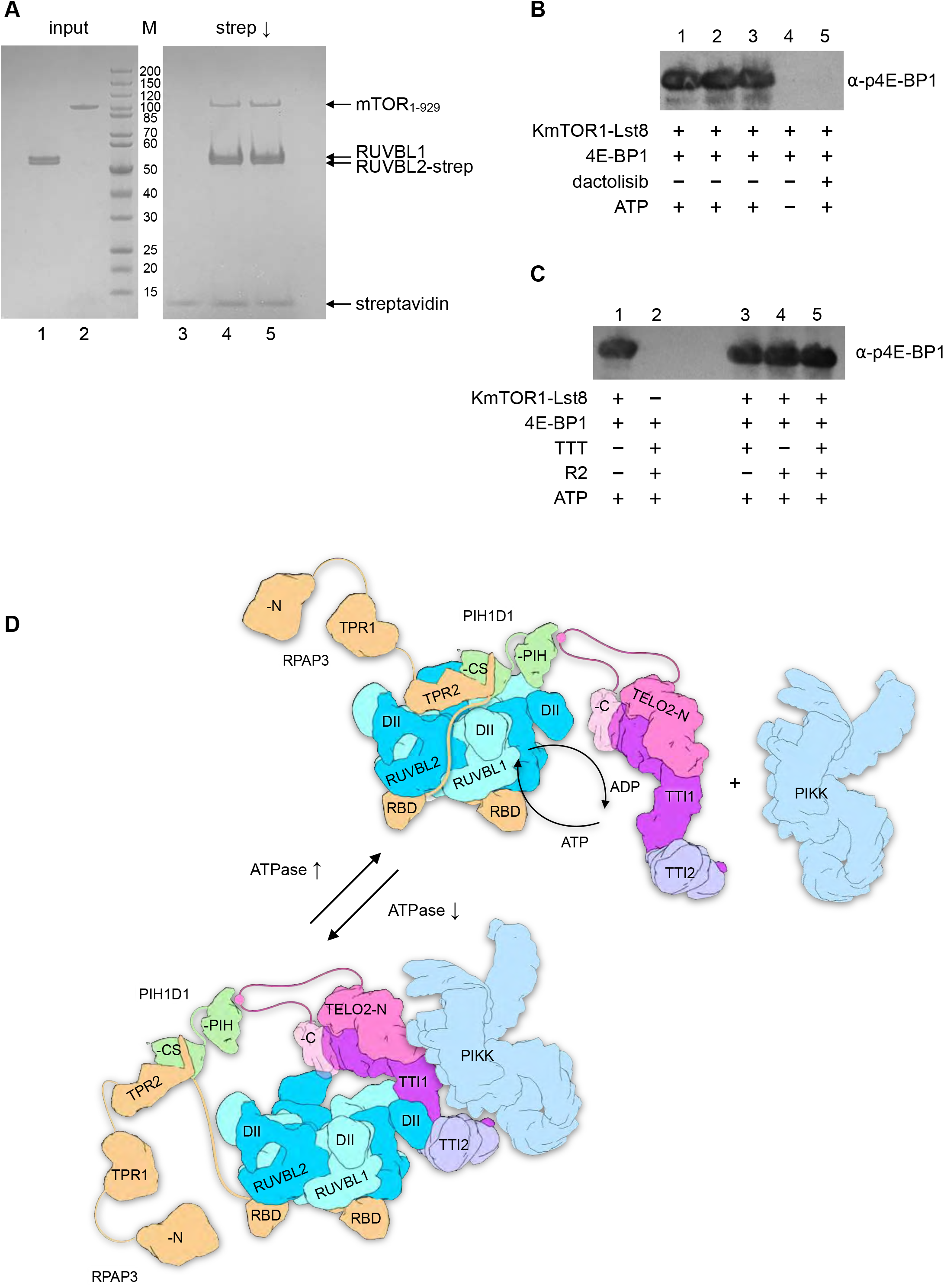
TTT couples mTOR recruitment to R2TP to regulation of the chaperone. **A.** Coomassie-stained SDS-PAGE gel showing the interaction of an mTOR fragment comprising the N-terminal HEAT region (residues 1-929) with strep-tagged RUVBL1-RUVBL2 complex. **B.** Western blot showing phosphorylation of 4E-BP1 by the *K.marxianus* TOR1-Lst8 complex. A strong signal was detected by a phospho-specific antibody to pThr37/46 in the presence of ATP, but not when ATP was omitted from the reaction, or when the ATP-competitive mTOR inhibitor dactolisib (NVP-BEZ235) was present. C. As B., but with the addition of TTT, R2, or both. Neither of these chaperone components, alone or in combination, affect its ability to phosphorylate a substrate. D. Model of the functions of TTT. Flexible tethering to the R2 ring, allows for facile exchange of TP and TTT components at the main interaction site on the DII-domain face of the ring. Interaction of the PIH1D1 part of TP with a RUVBL1 DII domain facilitates partial ring opening and nucleotide exchange (Munoz-Hernandez et al., 2019), accelerating the basal ATPase activity of the ring (Rivera-Calzada et al., 2017). The CK2-phosphorylation site within the inherently disordered linker connecting the N- and C-terminal domains of TELO2 (Horejsi et al., 2010) tethers the TTT sub-complex to the R2 ring through interaction with the PIH domain of PIH1D1 (Horejsi et al., 2014; Pal et al., 2014), and thereby facilitates recruitment of a PIKK client to the core complex (top). Rearrangement of the complex allows TTT to bind to the DII domains of consecutive RUVBL1-RUVBL2 domains, fixing them in an ADP-bound closed state that down regulates the basal ATPase, and potentially bringing the PIKK client into closer association. Steric hinderance of the DII domains and allosteric incompatibility, prevents simultaneous binding of PIH1D1 to the DII face of the ring. Although not engaged, TP remains tethered to the complex through the interaction of the PIH domain with the TELO2 linker segment (see above), and the interaction of the C-terminal domain of RPAP3 with the ATPase face of the R2 ring (bottom).

Taken together these data suggest that RUVBL1-RUVBL2 and TTT cooperate to bring TOR to the complex. The binding site for TTT on TOR actually resides in the C-terminal kinase region, rather than in the N-terminal HEAT repeat region, as had previously been suggested (Takai et al., 2007), and primarily involves a substantial direct contribution from TTI1/Tti1p-TTI2-Tti2p rather than TELO2/Tel2p itself (Takai et al., 2010). RUVBL1-RUVBL2 instead interacts with mTOR N-terminal HEAT repeat region.

Since TTT and R2 form direct complexes with mTOR, we sought to determine whether these interactions affected the ability of TOR to phosphorylate its substrates. We used a phosphospecific antibody to a highly conserved and well characterised TOR target phosphorylation site in 4E-BP1 (Wu et al., 2017) to show that our purified yeast KmTOR1-Lst8 was active as a kinase (Fig 5B), and was effectively inhibited by an ATP-competitive mTOR inhibitor (Liu et al., 2009). However, we found that addition of an excess of TTT or R2 or both together, had no effect on the ability of KmTOR1 to phosphorylate 4E-BP1 (Fig 5C), suggesting that the interactions these components of the R2TTT complex make with TOR proteins, does not have a role in regulating TOR kinase activity directly.

## Discussion

The heterohexameric RUVBL1-RUVBL2 ring appears to provide the central constant component of a multiplicity of chaperone complexes each specific for a distinct class of client proteins (Houry et al., 2018; Kakihara and Houry, 2012; Munoz-Hernandez et al., 2018). As with systems such as HSP90, with which RUVBL1-RUVBL2 (R2) collaborates as a co-chaperone, client specificity is believed to be mediated by scaffold proteins that structurally couple the generalised core R2-ATPase complex to the client. In both yeast and mammalian systems, the TELO2-TTI1-TTI2 / Tel2p-Tti1p-Tti2p (TTT) complex, whose structure we describe here, behaves as a classical scaffold adaptor, being able to independently bind a PIKK protein and the RUVBL1-RUVBL2 ring (**Fig. 1, 4**).

Current models suggest that recruitment of TTT to R2 is mediated by the interaction of a conserved phosphopeptide motif within the intrinsically disordered linker segment of TELO2/Tel2p, which binds to the PIH domain of PIH1D1/Pih1p (Horejsi et al., 2014; Horejsi et al., 2010; Pal et al., 2014) in the TPR protein - PIH (TP) component of an R2TP-TTT super-complex. However, our structural and biochemical data clearly shows that human TTT is fully able to bind R2 in the absence of PIH1D1 and RPAP3 (**Fig. 1**), so while this interaction may contribute to the function of the system, it is not essential for TTT recruitment. In the human system, the TTT and TP sub-complexes are able to coexist simultaneously in a complex with the same R2 ring, as demonstrated by the clear presence of density for RPAP3-RBD domains on the ATPase face of the R2 ring to TTT, in cryoEM reconstructions (**Fig. 3A**). However, no density was evident that would correspond to the expected location, based on our previous structural analysis (Munoz-Hernandez et al., 2019), of PIH1D1 attached to one of those RPAP3 molecules and engaged with the DII side of the R2 ring. The absence of density for PIH1D1, nonetheless present in the complex through its constitutive association with RPAP3, suggests that binding of TTT and PIH1D1 to the DII face of the R2 ring, may be mutually exclusive.

In the low-resolution yeast cryo-EM data, we identify only two classes of particle, in which either TP or TTT sub-complexes, but not both, are bound to the DII face of the R2 ring (**Fig. 3C, D**). The absence of any substantial set of particles displaying density for both TP and TTT components, implies that as in the human system, binding of these components to the R2 ring is mutually exclusive.

Taken together, our observations for human and yeast systems suggest that TTT and TP components are alternative R2 ligands engaging with the DII face of the ring with mutual exclusivity. However, the independent pairwise interactions of the TTT, TP, and R2 components, keeps the non-engaged component tethered to the overall complex. In the yeast system, this tethering is provided by binding of the PIH domain of Pih1p in the TP component to a putative CK2 phosphorylation site within an intrinsically disordered segment of ~50 residues that links the N-terminal and C-terminal domains of Tel2p. In the human system, in addition to the equivalent interaction of the PIH1D1 PIH domain and the CK2 phosphorylation site in the longer ~70 residue TELO2 linker segment, the TP component is also tethered by the C-terminal RBD domain of RPAP3 which connects by an intrinsically disordered segment of ~100 residues to the binding motif of for the CS-domain (Henri et al., 2018) (**Fig. 5D**).

This flexible tethering mechanism might serve to allow the non-engaged component to nonetheless remain associated with the overall complex and poised to exchange with the engaged component as required. We previously showed that binding of the TP component accelerated the inherent ATPase of the R2 ring (Rivera-Calzada et al., 2017), probably by promoting ring opening to facilitate nucleotide exchange (Munoz-Hernandez et al., 2019). Here we show that binding of the TTT complex has the converse effect, decreasing ATP turnover by stabilising the ring in a closed ADP-bound state. The flexible exchange of TP and TTT sub-complexes might therefore function in regulating the ATPase activity of the system at different stages of loading, manipulation and release of a PIKK client protein, however further work will be required to test this.

TELO2/Tel2p and the associated TTI1/Tti1p-TTI2/Tti2p are implicated in the activation and stabilisation of most, if not all members of the PI3-kinase-like kinase family (Takai et al., 2007). These proteins share strongly conserved C-terminal kinase regions attached to more diverse N-terminal segments composed of HEAT repeats. Paradoxically, a previous cell-based interaction analysis using expressed fragments of mTOR or ATM, had implicated segments of the less conserved HEAT repeats in the interaction with TELO2 (Takai et al., 2007). Using purified and soluble segments of mTOR, we find instead that it is the strongly conserved kinase-containing C-terminal region that is recognised by the TTT complex. However, this interaction does not inhibit kinase activity, indicating that it neither allosterically regulates TOR nor prevents access of substrate to its active site (**Fig 5C**). While we cannot dismiss the possibility that parts of the HEAT repeats of particular PIKKs may contribute to their individual interaction with TTT, the primary involvement of the common and highly conserved kinase region, provides a more evolutionarily compelling explanation for the selectivity of the TTT component of this chaperone/co-chaperone system. Nonetheless, RUVBL1-RUVBL2 binds independently of TTT to the N-terminal HEAT regions in mTOR, suggesting that both TTT and R2 components collaborate to mediate PIKK interactions.

Together, our results reveal the architecture of the TTT complex, and show how TTT couples the R2TPs recruitment of PIKKs to the regulation of the chaperone’s ATPase activity. The detailed structural basis for the interaction of TTT and its complexes with PIKKs and their complexes, and the roles these play in their activation and stabilisation, remain to be determined.

## Resource Availability

### Lead contact

Further information and requests for resources and reagents should be direct to and will be fulfilled by the Lead Contact, LHP

### Materials availability

Plasmids and constructs generated in this study can be requested from the lead contact.

### Data and code availability

Real space-refined atomic coordinates for the R2-TTT model have been deposited in the Protein Databank (PDB)with accessions code: ???. The composite cryoEM volume for R2-TTT has been deposited in the Electron Microcopy Databank (EMDB) with accession code: ???

### Experimental Model and Subject Details

#### Methods Details

The bacterial strain BL21 (DE3) *pLysS* was used to express yeast Tel2p proteins. The human and yeast TTT (Tel2-TTI1-TTI2) protein complexes, and human mTOR^N^ and mTOR^ΔN^-Lst8 were expressed in Sf9 cells. The yeast *Kluveromyces marxianus* yeast was used to express full-length KmTOR1-Lst8 complex.

#### Gene cloning and protein expression and purification

Yeast (*Saccharomyces cerevisiae*) and human R2 (RUVBL1/2 or Rvb1/2) and TP (RPAP3-PIH1D1 or Tah1p-Pih1p) were expressed and purified as previously described (Martino et al., 2018; Rivera-Calzada et al., 2017). *Kluveromyces marxianus* KmTOR-KmLst8 were purified as described in (Baretic et al., 2016). Human and yeast TELO2/Tel2p were cloned into pFBDM plasmid as BamHI and EcoRI fragments, respectively, in fusion with a GST-tag. The yeast Tel2p was also cloned into p3E (Sussex University) using NdeI and BamHI fragment resulting into GST-Tel2p which was co-overexpressed with CKII (Sussex University) in BL21 (DE3) *pLysS* cells (Thermo Fisher Scientific). TTI1 and GST-TTI2 were cloned into pFBDM plamid as XhoI-KpnI and BamHI-EcoRI fragments, respectively (Genscript). Strep-TTI1_1-459_ and Strep-TTI1_468-1089_ were cloned into pFBDM-GST-TTI2 plasmid for coexpression. GST-TTI2_1-193_ and GST-TTI2_194_._508_ and GST-mTOR^N^ (residues 1-929) were cloned into BamHI and EcoRI in pFBDM-TTI1 plasmid for *Spodoptera frugiperda* (Sf9) expression. The mTOR^ΔN^ (residues 1376 to 2549) containing 6XHis-2XFlag-PreScission cleavable tag at its 5’ was purchased from Genscript and cloned into BamHI and EcoRI sites of pFBDM and 5’ 2XFlag-6XHis-PreScission cleavable tagged-mLst8 gene was cloned as XhoI and KpnI fragment into pFBDM plasmid. TTI1-TTI2, full length as well as truncated versions and mTOR^ΔN^, mTOR^N^ complexes were overexpressed in Sf9 cells. For the co-expression of the yeast and human TTT complex, mixed viruses of yeast and human TTI1-TTI2/Tti1p-Tti2p and TELO2/Tel2p were used to co-infect Sf9 cells. Cell pellets containing yeast Tel2p were disrupted by sonication, while those containing TTI1-2 complex were lysed using a cell homogenizer in 20 mM HEPES pH 7.5, 250 mM NaCl,0.5mM TCEP (HEPES buffer) and 1 tablet of EDTA free protease inhibitors (Roche, 11697498001). The cell lysate was centrifuged at 21,000 g for an hour 4°C. The clear supernatant was added to equilibrated GST beads in HEPES buffer for affinity chromatography. The proteins were eluted using 50mM reduced glutathione from the GST beads. The GST-tag was cleaved with PreScission protease at 4°C overnight. The quality of the protein complexes was analysed using SDS-PAGE and the protein complexes were further purified by gel filtration using Superdex 200 increase 10/300 GL (GE healthcare) and Superose 6 10/300 (GE healthcare) column equilibrated in the HEPES buffer.

#### Purification of mTOR-KD-Lst8 complex and mTOR1-929 (heat repeat)

Human mTOR^ΔN^ (residues 1376 to 2549 with a His-Flag-PreScission cleavable tag at the N-terminus) was co-expressed with full length human Lst8 in *Spodoptera frugiperda* Sf9 cells. Cells were harvested and resuspended in lysis buffer containing 20 mM HEPES, pH 7.85, 250 mM NaCl, 1 mM EDTA, 0.5 mM TCEP and supplemented with 1 tablet of EDTA free protease inhibitors (Roche, 11697498001).Cells were lysed, centrifuged at 21,000 g for 60 min at 4°C and then applied to 1 ml of anti-FLAG M2 Affinity Gel-resin (Sigma-Aldrich Ltd) and incubated for 60 min at 4°C on a 200 rpm/min roller. The resin was then washed with 50 mL lysis buffer. The bound mTOR^ΔN^ was released from the resin using 0.2 mg/ml flag peptide (Peptide synthetics) dissolved in lysis buffer.The flag-tag was removed from mTOR-KD by incubating with PreScission protease at 4°C overnight. The mTOR^ΔN^ was then concentrated and further purified by gel-filtration using Superdex 200 increase 10/300 GL column in lysis buffer.

#### Interaction studies using pull down experiments

For GST-tag affinity, strep-tag and flag-tag pull-down experiments, 40μl beads were equilibrated in 20mM HEPES, pH 7.85, 140mM NaCl, 0.5mM TCEP, 0.03% CHAPS detergent. 20μl of 1mg/ml of GST-TTI2-TTI1, yeast and human TTT proteins, 1mg/ml C-terminal Strep-tagged RUVBL2 and 2XFlag-mTOR^N^, KmTOR-KmLst8 were used for pulldown experiments. The subcomplexes were incubated with either 40ul GST/Strep/Flag beads and incubated for 45mins at 4°C rotating at 20 r.p.m. The beads were washed three times with 200μl buffer by pelleting with quick centrifuge at 3000 g for 10sec. The quality of the pull-down interactions was visualised using 4-12% SDS-PAGE in MES and MOPS buffer.

#### ATPase assay

The ATPase assay of RUVBL1-RUVBL2 was carried using PiColorLock detection reagent kit (Innova Biosciences Ltd). The ATPase assay was carried out in 50mM Tris-HCl Ph 7.5, 50mM NaCl, 20mM MgCl2 0.1%Glycerol, 0.01% Tween X-100. All the ATPase reactions were carried out in triplicates. For each ATPase reaction, 0.25μM RUVBL1-RUVBL2, and 1.2μM-400μM ATP was used. The rest of the protocol was followed according to manufacturer’s instructions (Innova Biosciences Ltd). For the R2-TTT ATPase assay, 0.25μM RUVBL1-RUVBL2 and 0.75μM TTT complexes were mixed with above mentioned buffer and the PiColorLock reagents. The reaction mixture was incubated at 37°C for 30mins. For the negative control, 400 μM ATP alone and/or proteins alone were used. The reactions were scanned at 650nm using CLARIOstar (Biomedical Solutions Inc.).

#### TOR kinase assay

To measure the effect of TTT and R2TP in mTOR kinase activity, we used 1μM KmTOR-Lst8 1μM, 3μM Rvb1p/2p (hexamer) and 5.5μM ScTel2-TTi1-TTi2, and 16μM purified 4EBP1 protein. The kinase assay was performed using HEPES buffer (25mM HEPES, pH 7.5, 50mM KCl, 5mM MnCl2, 5mM MgCl2 in a final volume of 20μl. The kinase reaction was conducted in the presence of 100μM ATP at 23 □ for 30mins. For the negative control experiment, we used 10μM ATP competitive mTOR kinase inhibitor Dactolisib (company). The reaction was stopped by the addition of 20mM EDTA followed by the addition of 5μl Laemmli buffer followed by immediate boiling for 5mins at 100□. The samples were subjected to standard western blot using 4-12% SDS-PAGE. The phosphorylated 4EBP1 was detected by using anti-Phospho-4EBP1 (Thr37/46) polyclonal antibody (Cell Signalling Technology, cat. number 236B4). Chemiluminescence reaction was visualised using ECL solution (GE Healthcare) according to the manufacturer’s instructions.

#### Reconstitution of R2TP-TTT and R2-TTT complexes

1μM of the human and yeast R2TP/RUVBL1-RUVBL2 complexes was assembled according to previously described method (Martino et al., 2018; Rivera-Calzada et al., 2017). 2μM of pure yeast and human TTT complex were mixed with R2TP/RUVBL1-RUVBL2 complexes and these assembled complexes were incubated for 1 hour at 4°C. The quality of the samples was analysed using 4-12% SDS-PAGE and ultimately used for the cryo-grids preparation.

#### Cryo-EM Sample Preparation and Data Collection

Freshly assembled preparations of the different complexes (1μM) were used to prepare cryogrids by applying 3.7 μl of complex to a holey-carbon grids (Quantifoil™ 1.2/1.3, 300 mesh copper) which was glow discharged in air with PELCO easiGlow by applying a current of 10mA for 60 seconds. Grids were blotted for 3 seconds at 25°C and plunged into liquid ethane using FEI-Vitrobot Mark IV (Thermo Fisher Scientific) or Leica EM GP2 (Leica microsystems) and stored in liquid nitrogen prior to imaging. Data were collected using FEI Titan Krios electron microscopes at 300kV, equipped with a Gatan BioQuantum (slit width 20eV) K2 direct electron detectors operating in counting mode. The R2-TTT and R2-T2 (TTi1-TTi2) data were collected at Astbury Biostructure Laboratory, Leeds, UK and R2TP-TTT data was acquired at eBIC Diamond Light Source, UK. Automated data acquisition software EPU (Thermo Fisher Scientific) was used to collect data based on a published protocol (*29*) and with the parameters described in **Table S1**. Micrograph movies were collected with 40 fractions across a 10 s exposure. Autofocusing was performed after every 10 μm. For R2-T2 and R2-TTT data was collected using total doses of 49.6 e^-^/ Å ^2^ and 51.2 e^-^/ Å ^2^ respectively and the R2TP-TTT data was acquired with dose of 42.12 e/ Å ^2^ over 40 fractions.

#### Cryo-EM Data Processing

##### Cryo-EM Data Processing

All data sets (**Table S1**) were processed following similar strategies. Movies obtained for the different samples and imaging sessions were aligned and local motions were corrected using MotionCor2 (Zheng et al., 2017). The contrast transfer function (CTF) estimation was performed using Gctf (Zhang, 2016). Particle selection, 2D and 3D classifications, refinement and all other image processing steps were performed using the tools provided in Relion 2 (Kimanius et al., 2016) and Relion 3 (Zivanov et al., 2018), unless other programme is specified. Particles were automatically picked from the micrographs and the best particles in the initial data sets were selected after several rounds of 2D classification and averaging. Cleaned data sets were then classified in several sub-groups in 3D using initial templates generated using the *ab initio* reconstruction tools in cryoSPARC (Punjani et al., 2017) and the *ab-initio* routine available in Relion. The best sub-groups in each 3D classification were selected for refinement either separately or after merging those classes with similar features. The final structure was then selected and the particles were polished in Relion3.1 before a final round of refinement using shiny particles. In most maps, the resolution of the TTT complex was lower than the RUVBL1-RUVBL2 hexamer and we therefore implemented a multi-body refinement strategy (Nakane et al., 2018). We defined two bodies, corresponding to the TTT complex and the two DII domains in contact with TTT (body 1) and the RUVBL1-RUVBL2 hexamer except the two DII domains in contact with TTT (body 2). Multi-body refinement revealed some flexibility in the interaction between TTT and RUVBL1-RUVBL2 and significantly improved the resolution of the density corresponding to TTT. All final maps were post-processed and sharpened in Relion and the resolutions estimated in Relion using gold standard Fourier Shell Correlation (FSC) and a cut-off of 0.143 as implemented in Relion. Local resolution ranges for each map were estimated using Relion. Details about the initial and final number of images, estimated average resolution and range of resolutions for each of the maps described in this work are indicated in **Table S1**.

For the structure of the R2-TTT complex we saturated R2 with an excess of TTT and collected two independent data sets in two sessions using the same microscope and imaging conditions. The two datasets were initially processed separately and later merged as detailed in **Supplementary Figure 2** Under these saturated conditions, several TTT complexes interacted with each RUVBL1-RUVBL2 hexamer and we implemented a symmetry expansion strategy to make use of all the available information as described before (Martino et al., 2018; Zhou et al., 2015) Briefly, because each R2-TTT complex has a roughly threefold symmetry and contained one, two or three TTT complexes per R2, we rotated each particle around the 3-fold symmetry axis three times to place all TTT complexes in the same position. This operation triplicated the data set and then we placed a mask around one of the TTT positions and including part of the R2 ring. Particles were then subjected to a local classification strategy to select the best quality and more homogenous TTT complexes, which was then refined locally.

#### Model Generation, Fitting and Visualization

Atomic models for human RUVBL1 and RUVBL2 and RPAP3-RBD, were derived from previous cryoEM and NMR structures (Martino et al., 2018; Maurizy et al., 2018; Munoz-Hernandez et al., 2019) and initially docked as rigid bodies into cryoEM density in Chimera (Goddard et al., 2007). Homology models of the DII domains, which were not fully resolved in previous structures were constructed using I-TASSER (Roy et al., 2010), merged into the experimentally determined RUVBL1 and RUVBL2 models, docked into density using Chimera, and geometrically regularised in Coot (Casanal et al., 2019). A homology model for human TELO2 N-domain was constructed using the I-TASSER server. Side-chains were stripped back to Cβ except for proline residues, and the model docked as a rigid body into density using Chimera. The position and orientation of individual helices, and their connecting loops was manually optimised to the experimental density in Coot. Models for TTI1 and TTI2 were constructed by manual building of poly-alanine α-helices into experimental density, using the right-handed solenoid topology of HEAT repeat to determine connections. The polarity of the two chains was subsequently determined by biochemical analyses and the models adjusted accordingly. Based on the length of these proteins, the current TTI1 and TTI2 HEAT-repeat models account for ~60% and 90% of the amino acid sequences of these proteins, respectively. The fit of the overall model was optimised by real-space refinement in Phenix (Afonine et al., 2018) to a combined map generated from a consensus of the highest resolution overall map, and individual local-resolution filtered multibody-refined volumes (see above). All structural depictions were generated using Chimera.

#### Cross-linking mass spectrometry (XL-MS) of human R2TP-TTT complex

The 1uM assembled complex of R2TP-TTT (in 20 mM HEPES pH 8.0, 140mM NaCl, 0.5mM TCEP) was cross-linked with a 100-fold excess of isotopically labelled N-hydroxysuccinimide (NHS) ester disuccinimidyl suberate (DSS-H_12_/D_12_, Creative Molecules Inc., Canada), with respect to the protein concentration. The cross-linking reactions were incubated for 60 minutes at room temperature and then quenched by the addition of NH_4_HCO_3_ to a final concentration of 20 mM and incubated for further 15 min.

The cross-linked proteins were reduced with 10 mM DTT and alkylated with 50 mM iodoacetamide. Following alkylation, the proteins were digested with trypsin (Promega, UK) at an enzyme-to-substrate ratio of 1:50 in an overnight incubation at 37 °C followed by the addition of Glu-C protease (Promega, UK) at a ratio of 1:10 and incubated for 4 hours at 37°C. After digestion, the samples were acidified with formic acid to a final concentration of 2% v/v and the peptides fractionated by peptide size exclusion chromatography, using a Superdex Peptide 3.2/300 (GE Healthcare) with 30% v/v Acetonitrile/0.1% v/v TFA as mobile phase and at a flow rate of 50 μl/min. Fractions were collected every 2 min over the elution volume 1.0 ml to 1.7 ml. Prior to LC-MS/MS analysis fractions were freeze dried.

Lyophilized peptides for LC-MS/MS were resuspended in 0.1 % (v/v) formic acid and 2 % (v/v) acetonitrile and analyzed by nano-scale capillary LC-MS/MS using an Ultimate U3000 HPLC (ThermoScientific Dionex, USA) to deliver a flow of approximately 300 nl/min. A C18 Acclaim PepMap100 5 μm, 100 μm□×□20 mm nanoViper (ThermoScientific Dionex, USA), trapped the peptides before separation on a C18 Acclaim PepMap100 3 μm, 75 μm□×□250 mm nanoViper (ThermoScientific Dionex, USA). Peptides were eluted with a gradient of acetonitrile. The analytical column outlet was directly interfaced via a nano-flow electrospray ionisation source, with a hybrid dual pressure linear ion trap mass spectrometer (Orbitrap Velos, ThermoScientific, San Jose, USA). Data dependent analysis was carried out, using a resolution of 30,000 for the full MS spectrum, followed by ten MS/MS spectra in the linear ion trap. MS spectra were collected over a m/z range of 300–2000. MS/MS scans were collected using threshold energy of 35 for collision-induced dissociation.

For data analysis, Xcalibur raw files were converted into the MGF format through MSConvert (Proteowizard; (Kessner et al., 2008)) and used directly as input files for StavroX (Gotze et al., 2015). Searches were performed against an ad hoc protein database containing the sequences of the complex and a set of randomized decoy sequences generated by the software. The following parameters were set for the searches: maximum number of missed cleavages 3; targeted residues K, S, Y and T; minimum peptide length 5 amino acids; variable modifications: carbamidomethyl-Cys (mass shift 57.02146 Da), Met-oxidation (mass shift 15.99491 Da); DSS cross-links mass shift 138.06808 Da (precision: 10 ppm MS1 and 0.8 Da MS2); False Discovery Rate cut-off: 5%. Finally, each fragmentation spectrum was manually inspected and validated.

### Quantitation and Statistical Analysis

#### ATPase activity

Data were fitted to the Michaelis–Menten equation using non-linear regression in GraphPad Prism Software 8 to calculate the reported Vmax and Km values.

#### Key Resources Table

| REAGENT or RESOURCE | SOURCE | IDENTIFIER |
| --- | --- | --- |
| Bacterial and Virus Strains |  | Catalogue number |
| *E.coli* BL21 (DE3) *pLysS* | Thermo Fisher Scientific | 10604973 |
| *E.coli* DH5α | Thermo Fisher Scientific | 11583117 |
| *E.coli* DH10Bac | Thermo Fisher Scientific | 10361-012 |
| **Chemicals, Peptides, and Recombinant Proteins** |  |  |
| DSS-H<sub>12</sub>/D<sub>12</sub> | Creative Molecules Inc | 001S |
| Trypsin | Promega, UK | V5111 |
| Glu-C protease | Promega, UK | V1651 |
| 2XFlag-peptide | Peptide synthetics, UK | Quote Q23477 |
| L-Glutathione reduced | Thermo Fisher Scientific | 10628343 |
| ANTI-FLAG<sup>®</sup> M2 Affinity Gel | Sigma-Aldrich | A2220 |
| Complete, EDTA-free protease inhibitor tablets | Roche Diagnostics | 05056489001 |
| **PhosSTOP** | Sigma-Aldrich | 4906845001 |
| PreScission and TEV protease | University of Sussex | N/A |
| Phospho-4EBP1 (Thr37/46) polyclonal antibody , | Cell Signalling Technology | cat. number 236B4 |
| **Critical Commercial Assays** |  |  |
| P<sub>i</sub>ColorLock Gold Phosphate Detection System | Innova Biosciences | 303-0030 |
| Cellfectin II Reagent | Thermo Fisher Scientific | 10362100 |
| **Deposited Data** |  |  |
| **Experimental Models: Cell Lines** |  |  |
| Sf9 cells | Thermo Fisher Scientific | Cat# 11496015 |
| **Recombinant DNA** |  |  |
| P3E plasmid | University of Sussex | N/A |
| pFBDM plasmid expressing human and yeast TTT and mTOR-KD proteins | This study | N/A |
| KmTOR-KmLst8 | A Gift from Prof. Roger Williams | N/A |
| **Software and Algorithms** |  |  |
| PRISM 8.0 | GraphPad | https://www.graphpad.com/scientific-software/prism/ |
| Relion-3.1 |  |  |
| EPU software | Thermo Fisher Scientific | https://www.fei.com/software/epu-automated-single-particles-software-for-life-sciences/ |
| MotionCor2 | Zheng *et al.*, 2017 | https://msg.ucsf.edu/software |
| GCTF and CTFFIMND4 | Zhang, 2016<br>Rohou and Grigorieff, 2015 | https://www.mrc-lmb.cam.ac.uk/kzhang/Gctf/Gctf_v1.18_b2/<br>https://grigoriefflab.umassmed.edu/software_download |
| Phenix | Adams et al., 2010 | https://www.phenix-online.org/ |
| Coot | Emsley et al., 2010 | https://www2.mrc-lmb.cam.ac.uk/personal/pemsley/coot |
| Pymol | Schrödinger | https://pymol.org/2/ |
| Chimera | Petterson et al., 2004 | http://www.rbvi.ucsf.edu/chimera |
| I-TASSER | Yang et al., 2015 | https://zhanglab.ccmb.med.umich.edu/I-TASSER/ |
| Phyre<sup>2</sup> | Kelly et al., 2015 | http://www.sbg.bio.ic.ac.uk/~phyre2/html/page.cgi?id=index |
| **Other** |  |  |
| Quantifoil 1.2/1.3, 300 mesh copper | Agar Scientific | AGS143-2 |
| FEI-Vitrobot MAG IV | Thermo Fisher Scientific | N/A |
| Leica EM GP2 | Leica microsystems | N/A |
| Titan Krios | Thermo Fisher Scientific | N/A |
| K2 direct detector camera | GATAN | N/A |

## Supplementary Materials

Table S1. Cryo-EM Data Processing and Refinement Statistic

Fig S1. R2-TTT Complex Formation

Fig S2. R2-TTT Cryo-EM Data Processing

Fig S3. Cryo-EM of R2-TT and mapping TELO2

Fig S4. R2TP and TTT interactions

Fig S5. R2TP-TTT Cryo-EM Data Processing

Fig S6. TTT and TOR Interactions

## Acknowledgments

**General**

We are grateful to Pascale Schellenberger (University of Sussex), and Fabienne Beuron and Ed Morris (Institute of Cancer Research) for assistance with cryoEM grid preparation and evaluation, and for useful discussion. We are grateful for access to the cryoEM facility at the University of Sussex (funded by Wellcome Trust Award Enhancement Grant 095605/Z/11/A – to LHP - and the RM Phillips Trust), and FEI Krios Titan cryoelectron microscopes at the Astbury Biostructure Laboratory at the University of Leeds (funded by the University of Leeds ABSL award and Wellcome Trust award 108466/Z/15/Z) and at the UK national electron bioimaging centre (eBIC - Diamond, funded by the Wellcome Trust, MRC and BBSRC).

## Funding

This work was supported by a Welcome Trust Senior Investigator Award (095605/Z/11/Z) and Award Enhancement Grant (095605/Z/11/A) (LHP), a BBSRC Project Grant BB/R01678X/1 (LHP and CP), and by grants from the Spanish Ministry of Science and *Innovation/Agencia Estatal de Investigación* (MCI/AEI) co-funded by the European Regional Development Fund (ERDF)(SAF2017-82632-P), the Autonomous Region of Madrid co-funded by the European Social Fund and the European Regional Development Fund (Y2018/BIO4747 and P2018/NMT4443) and the support of the National Institute of Health Carlos III to CNIO (OL).

## Author Contributions

Conceptualization: M.P., L.H.P., O.L., C.P.; Methodology: M.P., L.H.P., O.L., C.P., J.M.S., R.F.T.; E.L.H Validation: J.M.S., R.F.T., L.H.P., O.L.; Formal Analysis: M.P., L.H.P., O.L.; Investigation: All Authors; Writing – Original Draft: L.H.P., O.L.; Writing – Review & Editing: M.P., L.H.P., O.L.; Visualisation M.P., L.H.P., O.L.; Supervision: L.H.P., O.L., C.P., J.M.S., R.F.T.; Funding Acquisition: L.H.P., O.L., C.P.

## Competing Interests

The authors declare no competing interests.

## Notes

### Competing Interest Statement

The authors have declared no competing interest.

### Summary of Updates

Corrected Author order Corrected Figure references

