## Supplementary material for "Structure of the TELO2-TTI1-TTI2 complex and its function in TOR recruitment to the R2TP chaperone"

**Table S1**: **Cryo-EM data collection, refinement and validation statistics**

|  | R2TP-TTT | R2-TTT  Data set 1 | R2-TTT  Data set 2 | TTT from R2-TTT | R2-TT | yR2TP-TTT |
| --- | --- | --- | --- | --- | --- | --- |
| **Data collection and processing** |  |  |  |  |  |  |
| Magnification | 130,000 | 130,000 | 130,000 | 130,000 | 130,000 | 130,000 |
| Voltage (kV) | 300 | 300 | 300 | 300 | 300 | 300 |
| Electron exposure (e–/Å^2^) | 50 | 51.2 | 51.2 | 51.2 | 49.6 | 50.1 |
| Defocus range (μm) | 1.20-2.70 | 1.25-2.5 | 1.25-2.5 | 1.25-2.5 | 1.25-2.50 | 0.75-2.0 |
| Pixel size (Å) | 1.048 | 1.07 | 1.07 | 1.07 | 1.07 | 1.07 |
| Symmetry imposed | C1 | C1 | C1 | C1 | C1 | C1 |
| Initial particle images (no.) | 168882 | 207723 | 446016 | 446016 | 57102 | 231596 |
| Final particle images (no.) | 90575 | 142043 | 267149 | 88340 | 23518 | 87438 |
| Map resolution (Å)  FSC (0.143) | 6.1 | 4.3 | 3.41 | 5.02 | 9 | 5.8 |
| Map resolution range (Å) | 4.39 - 12 | 3.72-7.99 | 3.19 – 8.19 | 4.07 – 9.9 |  |  |
| **Refinement** |  | using Phenix combine-focused map (EMDB xxxx)(PDB xxxx) | | |  |  |
| Initial model used (PDB code) |  | 6FO1 | | |  |  |
| Model resolution (Å)  FSC (0.143) |  | 3.3 | | |  |  |
| Map sharpening *B* factor (Å^2^) |  | 110 | | |  |  |
| Model composition  Non-H atoms  Protein residues  Ligands |  | 27830  4120  6 | | |  |  |
| *B* factors (Å^2^)  Protein  Ligand |  | 160.3  42.4 | | |  |  |
| R.m.s. deviations  Bond lengths (Å)  Bond angles (°) |  | 0.007  0.899 | | |  |  |
| Validation  MolProbity score  Clashscore  Poor rotamers (%) |  | 3.1  78.2  0.23 | | |  |  |
| Ramachandran plot  Favored (%)  Allowed (%)  Disallowed (%) |  | 79.91  19.77  0.32 | | |  |  |

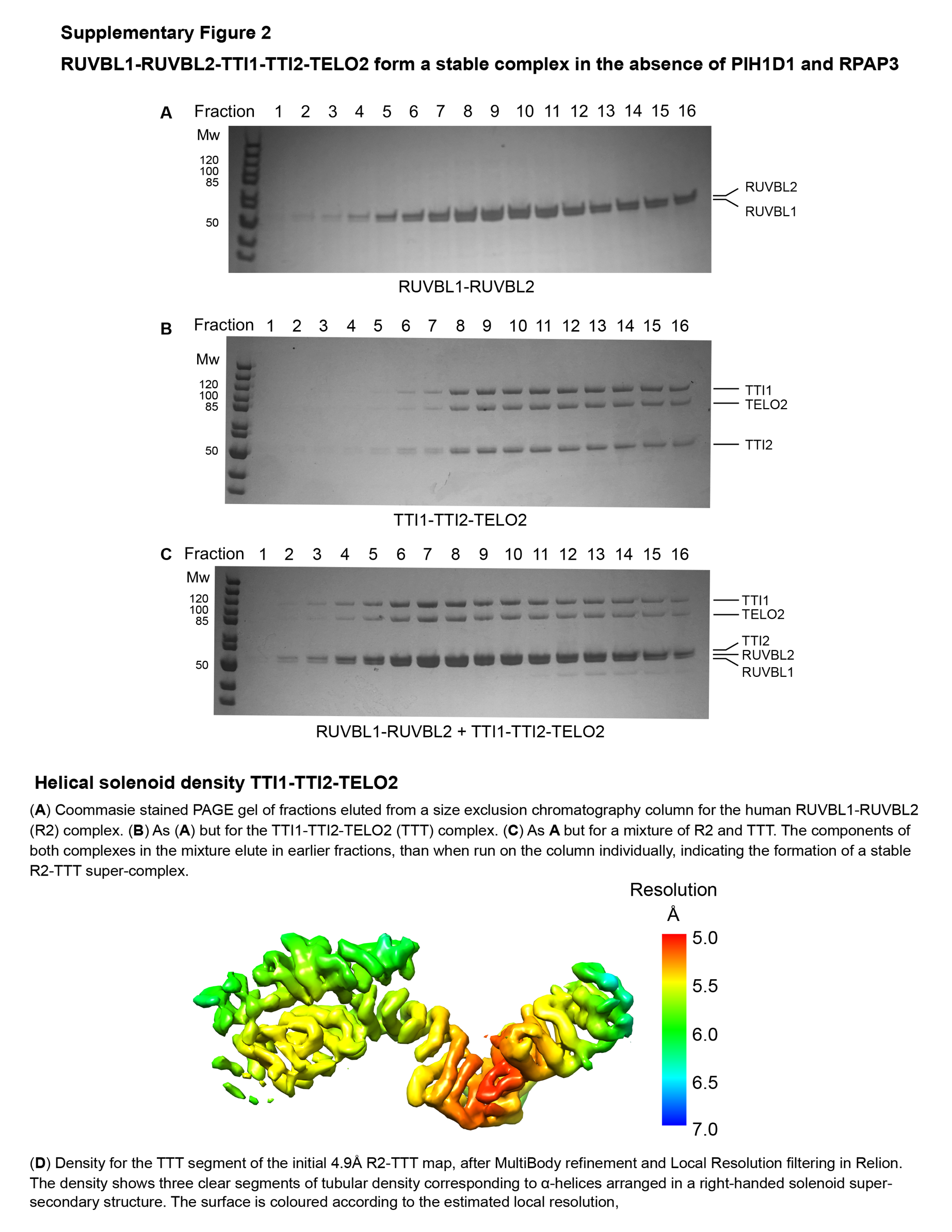

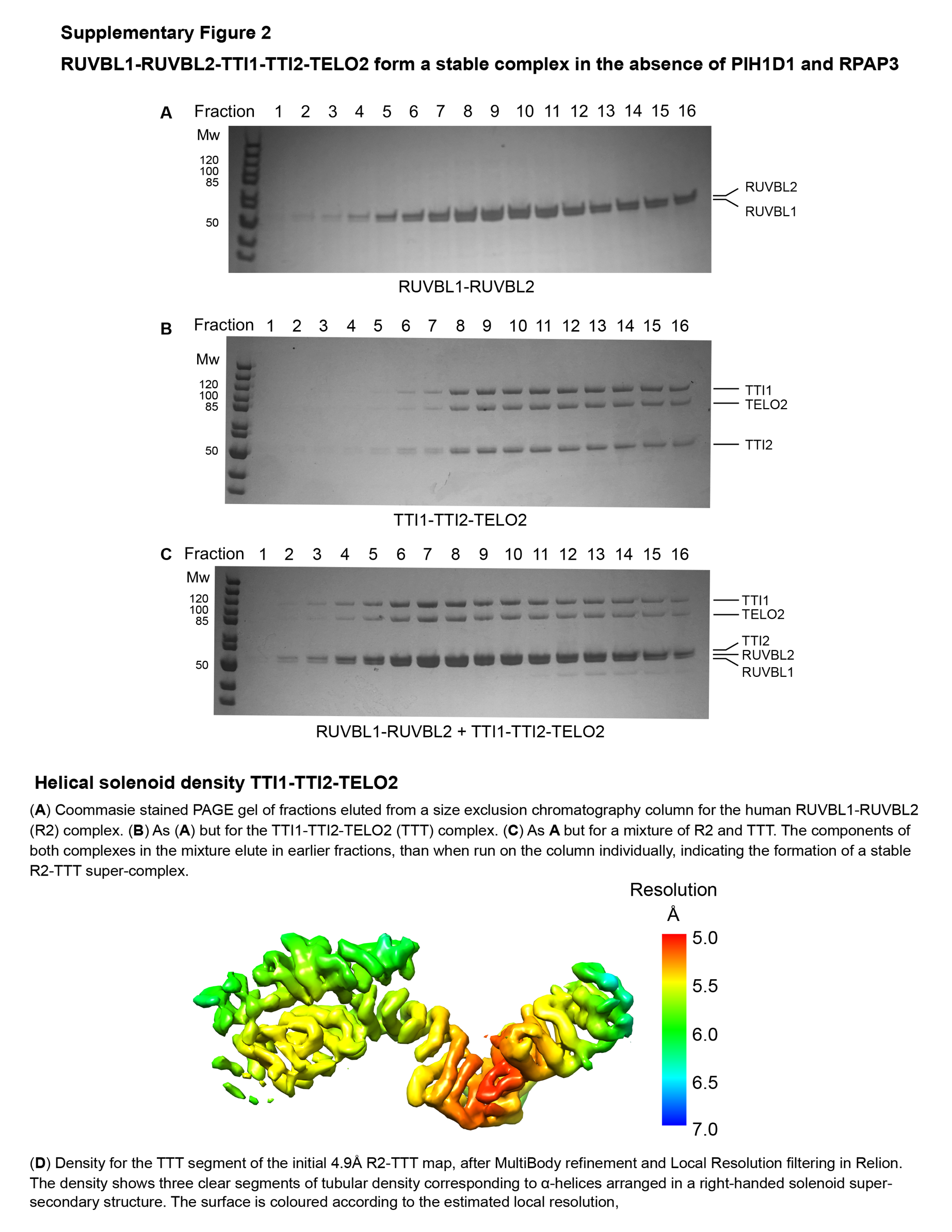

Supplementary Figure 1 – relates to Fig 1

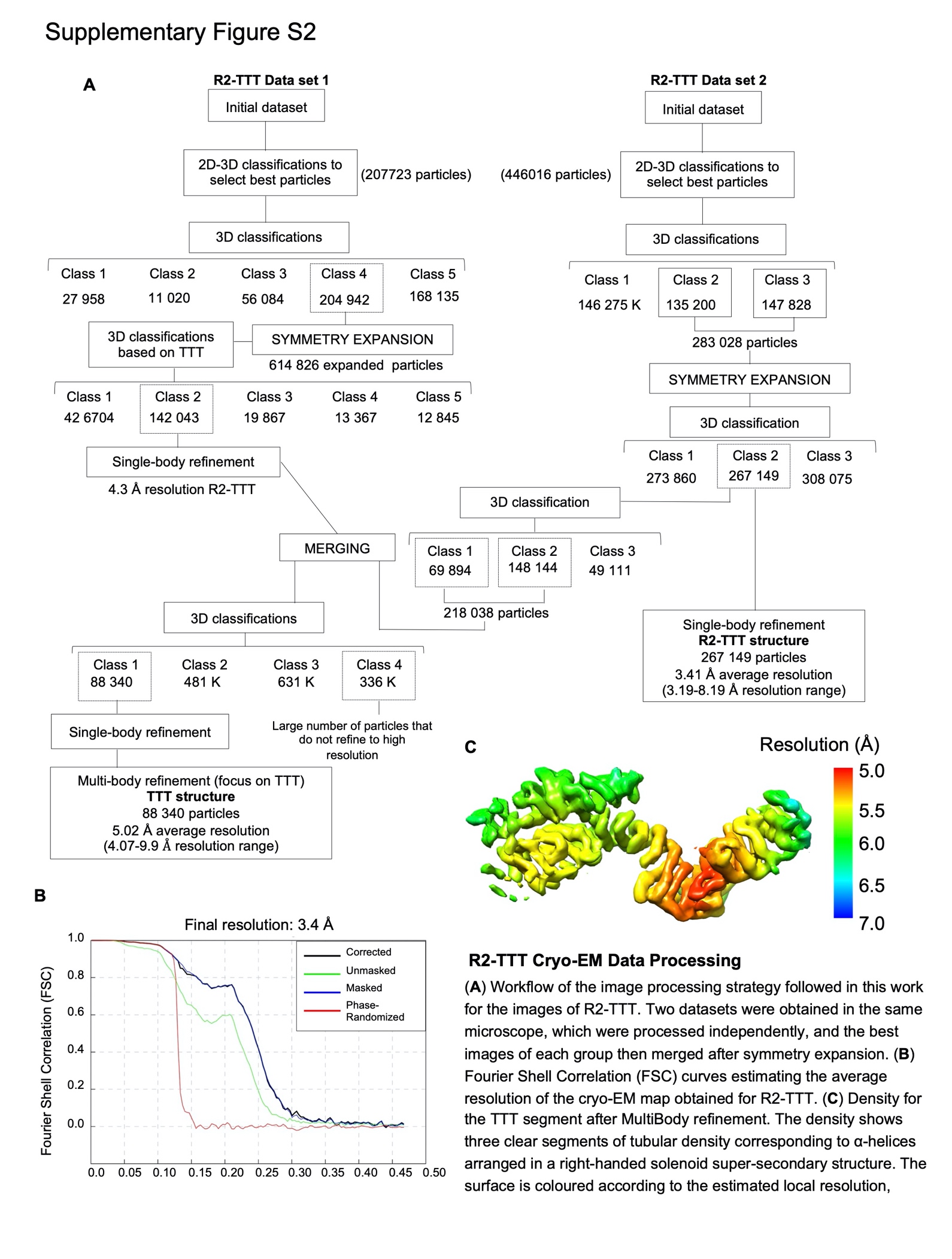

Supplementary Figure 2 – relates to Fig 1

Supplementary Figure 3– relates to Fig 1

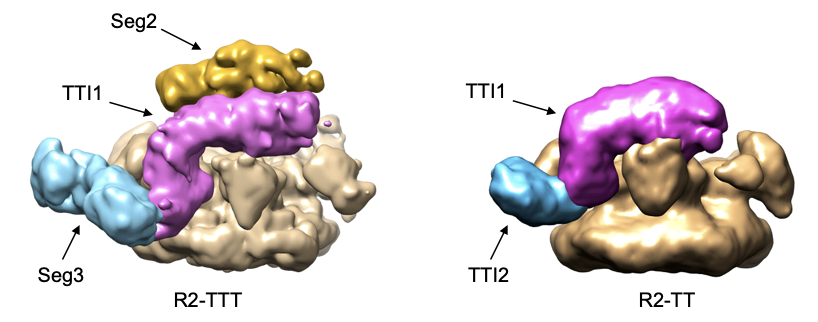

Comparison of cryoEM structure of (left) R2-TTT complex (5.4Å resolution) with (right) R2-TT complex (~9Å resolution), showing the absence of Seg2 (gold) in the R2-TT complex from which TELO2 was omitted. Both maps are shown filtered to 9Å to facilitate comparison.

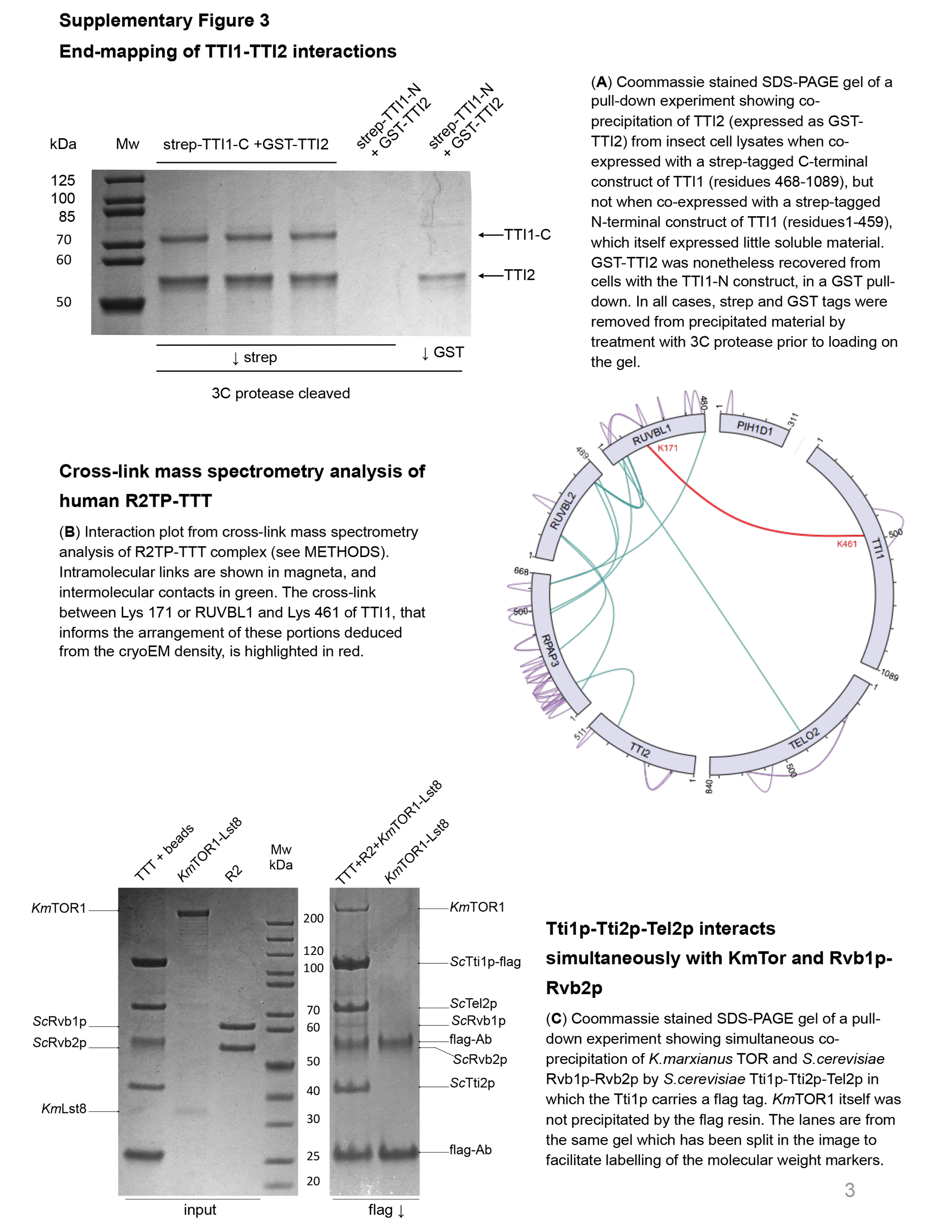

Supplementary Figure 4 – relates to Fig 2

Supplementary Figure 5– relates to Fig 3

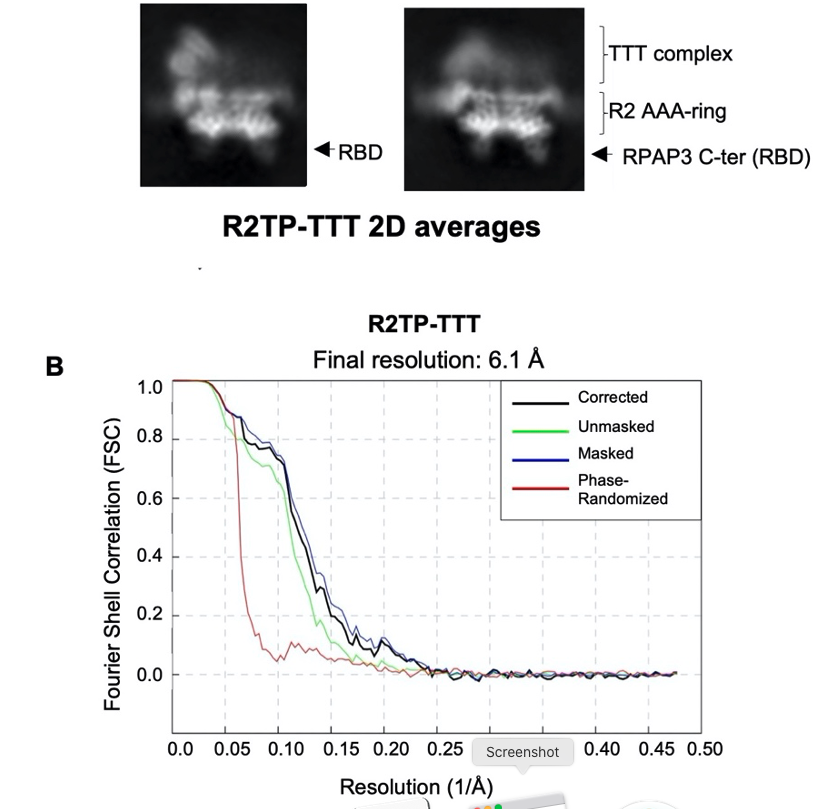

A. Representative 2D averages for R2TP-TTT, indicating the positions of the TTT comple, the AAA-ring of RUVBL1-RUVBL2 (R2) and the RBD domain of RPAP3
B. Fourier Shell Correlation (FSC) curves estimating the average resolution of the cryo-EM map obtained for R2TP-TTT

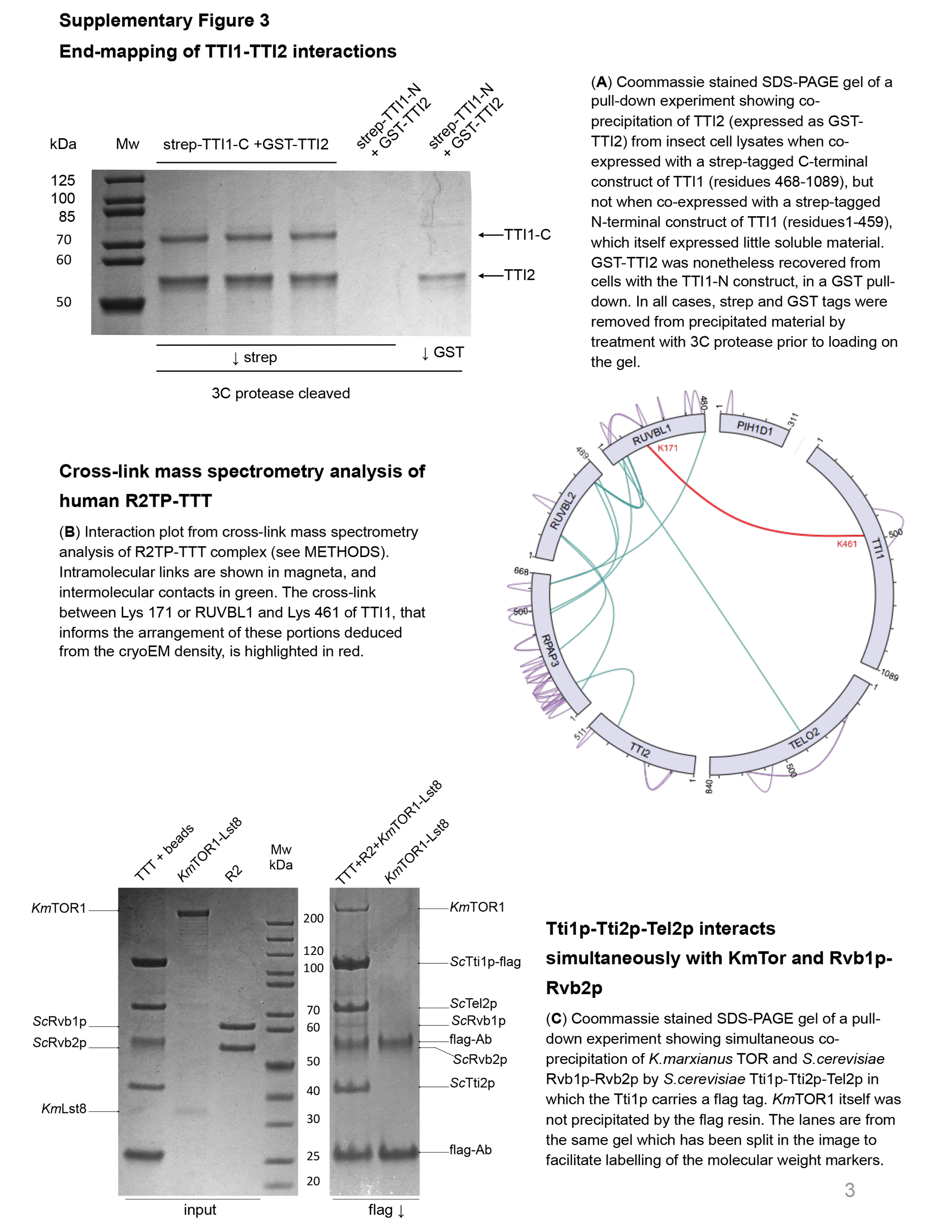

Supplementary Figure 6 – relates to Fig 4
